# Emergence of sympatric host-specific lineages of the fungal plant pathogen *Zymoseptoria passerinii* in natural ecosystems

**DOI:** 10.1101/2024.07.12.603051

**Authors:** Idalia C. Rojas-Barrera, Victor M. Flores-Núñez, Janine Haueisen, Alireza Alizadeh, Fatemeh Salimi, Eva H. Stukenbrock

**Affiliations:** Christian-Albrechts University of Kiel, Environmental Genomics, Am Botanischen Garten 1-11, 24118 Kiel, Germany; Max Planck Institute for Evolutionary Biology, August-Thienemann-Str. 2, 24306 Plön, Germany; Department of Plant Protection, Faculty of Agriculture, Azarbaijan Shahid Madani University, Tabriz, Iran; Department of Plant Protection, Faculty of Agriculture, University of Tehran, Karaj, Iran; Department of Biological Sciences, Institute of Ecology, Evolution and Diversity, Goethe University, Max-von-Laue Str. 13, D-60438, Frankfurt am Main, Germany

**Keywords:** Plant-pathogen phylogenetics, Pathogens in crop origin centers, Septoria speckled leaf blotch, Wild-plant pathogens, Wild pathosystems

## Abstract

- The barley disease Septoria Speckled Leaf Blotch, caused by the fungal pathogen *Zymoseptoria passerinii,* had its last outbreak in North America in the early 2000s. Although rare in agricultural settings, field sampling of wild grasses in the Middle East revealed the disease persistence in wild barley.
- Identification of *Z. passerinii* in distinct wild barley species led us to investigate signatures of host specialization using genomics to address the mode of emergence by host tracking or host range expansion. Furthermore, we applied virulence assays and confocal laser microscopy to evaluate if the disease development differs between wild and domesticated barley.
- Wild- and domesticated-host infecting populations have diverged, and phylogenetic relationships support the emergence of sympatric host-specific lineages. Cross-virulence assays showed that *Zymoseptoria* passerinii from domesticated hosts infect domesticated barley and its wild ancestor, *Hordeum spontaneum*. However, wild isolates from Iran did not infect domesticated barley. Wild and domesticated pathosystems have similar disease timing and progression, suggesting its persistence in natural ecosystems might be tied to environmental conditions.
- The study supports that a wide range of hosts can foster the emergence of host-specific lineages in sympatry and provide novel insights into the evolution of understudied fungal pathogens on wild crop relatives.

## Introduction

The increasing emergence and severity of infectious fungal diseases threaten food security and natural ecosystems (Fisher *et al*., 2012; Stukenbrock & Gurr, 2023). Continuous monitoring, prediction modeling of disease spread, and deeper comprehension of fungal pathogens in wildlife and wild plant hosts have been largely neglected. These are crucial to profile the impact of fungal pathogens in the context of climate change and independent of agricultural environments (Fisher *et al*., 2012). Current evidence supports that crop wild relatives (CWRs) might serve as reservoirs for domesticated plant pathogens (Monteil *et al*., 2013, 2016), although still few studies are focused on wild pathogen population processes and dynamics (Rouxel *et al*., 2013; Penczykowski *et al*., 2015; Eck *et al*., 2022; Treindl *et al*., 2023). CWRs hold higher levels of genetic diversity and have coevolved with plant pathogens in natural ecosystems. Moreover, the centers of diversity and domestication of crop plants harbor a wealth of species (Harlan, 1971) that could serve as hosts for plant pathogens. Despite the latter, natural ecosystems are undervalued economically, which limits funding for studies (Fisher *et al*., 2012). Furthermore, having access to wild species found in remote locations or immersed in complex geopolitical contexts adds another layer of difficulty, generating a geographical bias toward high-income regions at the expense of exploring the remaining biodiversity (Marks *et al*., 2023). One way to overcome this is to prioritize neglected areas by collaborating with scientific communities situated in less-represented regions of the globe (Marks *et al*., 2023), and promoting research on non-model species and dynamics in natural ecosystems.

Cumulative evidence supports that ecological divergence of plant pathogens is driven by host specialization. As proposed by Crous & Groenewald (2005) and exemplified by multiple studies (Steenkamp *et al*., 2002; Choi *et al*., 2011; Rouxel *et al*., 2013; Faticov *et al*., 2022), plant pathogens phylogenies frequently represent several closely related sister or cryptic species. In this regard, the *Zymoseptoria* genus comprises eight ascomycete species, only two of them, *Z. tritici* and *Z. passerinii*, have been reported to infect domesticated hosts (Quaedvlieg et al., 2011; Stukenbrock et al., 2012b). The origin, population genetics, and plant-pathogen dynamics of the prominent wheat fungal pathogen *Z. tritici* have been extensively investigated in an agricultural context (Linde *et al*., 2002; Stukenbrock *et al*., 2011; Orellana-Torrejon *et al*., 2022; McDonald *et al*., 2022; Feurtey *et al*., 2023), but despite *Z. tritici* and *Z. passerinii* having alternative wild hosts (See (Rojas-Barrera *et al*., 2023) for a summary), our understanding of the populations genetics and pathogens prevalence in natural ecosystems remains limited (Stukenbrock *et al*., 2011, 2012a). Population genetic studies on *Z. tritici* support that the center of diversity of the *Zymoseptoria* genus is located in the Middle East (Banke *et al*., 2004), in proximity to the Fertile Crescent recognized as a center of crop domestication (Harlan, 1971), where also several CWRs were found to be abundant and naturally distributed (Harlan & Zohary, 1966).

The hemibiotrophic fungus *Z. passerinii* is the causal agent of the disease Septoria Speckled Leaf Blotch (SSLB), a sporadic disease that became important during its last outbreak in the late 1990s and the first half of the 2000s decade, in the Upper Midwest of the USA and neighbor provinces in Canada (Toubia-Rahme & Steffenson, 2004). During that period, an extensive study with 309 isolates collected in North Dakota and Western Minnesota in 2003 and 2004, revealed a high genetic diversity, and a shallow population structure for *Z. passerinii* (Lee & Neate, 2007b); which is supported by an equilibrated frequency of both mating types (Lee & Neate, 2007a), suggesting sexual reproduction. However, the teleomorphic stage of this pathogen has only been reported under experimental conditions and has not been described in the field (Ware *et al*., 2006). Interestingly, despite the relevance of SSLB at the beginning of the 2000s decade, there are, to our knowledge, no current reports of SSLB outbreaks in North America or elsewhere, including the Middle East is which is known to be a center of diversity of *Zymoseptoria*.

The disease triangle states that favorable climatic variables are as important as plant characteristics in determining the severity of disease epidemics (Stevens, 1960). Thus, the sporadic nature of SSLB has been attributed to the requirement of more than 48 h of continuous moisture for spore germination (Green & Dickson, 1957) and a long period of incubation (16–19 days) (Koble *et al*., 1959; Cunfer, 2000), which supports dependence on environmental conditions for *Z. passerinii* infection, additionally, the development of resistant cultivars with durable resistance traits (Toubia-Rahme & Steffenson, 2004), has been related to the abrupt disappearance of SSLB during recent decades.

In contrast to agricultural environments, our sampling of wild grasses during 2018 and 2020 revealed the persistence of SSLB in multiple *Hordeum* sp. in Northwest Iran, which overlaps with the Fertile Crescent region, and the center of origin and diversity for fungal plant pathogens and their wild host species (Harlan & Zohary, 1966; Banke et al., 2004).

In this work, we aimed to answer the following questions: Have the barley-infecting lineages o*f Z. passerinii* arisen through host tracking, similar to *Z. tritici* (Stukenbrock *et al*., 2011) or through host range expansion? Given our observation of symptoms in what were identified as different *Hordeum* species, we speculate whether host specificity in sympatric populations has led to population divergence in the center of diversity of the *Zymoseptoria* genus. Lastly, we ponder whether the persistence of SSLB in natural ecosystems is linked to the distinct disease onset observed in wild-host infecting lineages compared to the domesticated-host. We address these questions using population genomic datasets of host-specific populations of *Z. passerinii*.

In addition to the aforementioned inquiries, we offer insights into *Z. passerinii* populations within a natural ecosystem situated at the center of origin for both the pathogen and its hosts. Furthermore, we established a pathosystem in barley and the CWR *H. murinum* ssp. *glaucum*. Barley (2n) offers an alternative to hexaploid wheat, to study *Zymoseptoria*-caused disease in cereals and investigating the role of candidate resistance traits within a diploid genome background.

## Material and methods

### Fungal isolation, DNA extraction, and sequencing

Leaves of wild *Hordeum* species exhibiting SSLB symptoms were collected, in Northwest Iran, in 2018 and 2020. *Z. passerinii* isolates were purified following the methods described for *Z. tritici* (Fagundes *et al*., 2020). We obtained 59 isolates from five different wild grass lesions collected at four locations. Fungal species identification was performed by amplifying ITS1 (White *et al*., 1990; Baldwin, 1992; Baldwin *et al*., 1995). In order to recognize potential clones, we performed a PCR based genotyping. We performed clonal genotype discrimination with three ISSR markers (<u>Table S1</u>), and selected 34 of the 59 isolates for further genomic DNA extraction with a Phenol-Chloroform protocol (Fagundes *et al*., 2020). Additionally, we included nine *Z. passerinii* isolates from domesticated barley *(H. vulgare*) and one isolate from a wild grass collected in the US (Zpa924) (<u>Table S</u>3). These additional ten isolates were collected in four different counties in the northern USA. A total of 42 isolates, along with two technical replicates, were submitted for whole genome sequencing (2x150) with the Illumina HiSeq3000 sequencing platform at the Max-Planck-Genome-Centre, Cologne, Germany (http://mpgc.mpipz.mpg.de).

Herbarium samples were disrupted using a Precellys® instrument for the plant host identification, and DNA extraction was performed with the NucleoSpin Plant II, Mini Kit Macherey Nagel (REF 740770.50), following the manufacturer’s instructions. The markers ITS2, matK, trnH-psbA, atpB-rbcL, and trnL-trnF (Sang *et al*., 1997; Nishikawa *et al*., 2002; Yu *et al*., 2011; Bieniek *et al*., 2015; Ganopoulos *et al*., 2017), were amplified by PCR and sequenced by Sanger sequencing at the Competence Centre for Genomic Analysis (CCGA) Kiel, Germany. The sequences were trimmed and blasted with Genius Prime 2022.2. Subsequently, these were aligned and concatenated with MEGA11 (Tamura *et al*., 2021) to build a maximum likelihood tree using RAxML on CIPRES (Miller *et al*., 2015).

We selected the Iranian isolate Zpa796, to generate a high quality genome assembly for the Iranian *Z. passerinii* populations, DNA extraction was carried out with a modified CTAB protocol (Allen *et al*., 2006) using overnight lyophilized fungal cells. This sample was used for long-read sequencing PacBio Sequel II. Sequencing was conducted at the Max Planck-Genome-Centre, Cologne, Germany (https://mpgc.mpipz.mpg.de/).

### RNA extraction and sequencing

Three biological replicates of Zpa796 were collected after 72h of growth in YMS (yeast extract, malt-extract, and sucrose) liquid media and kept at -80 °C until further processing. RNA isolation was carried out with the Direct-zol ™ RNA Miniprep Plus Kit (cat. 2070, Zymo Research) following the manufacturer’s guidelines. The quality control, RNA seq libraries including poly-A enrichment, and sequencing on the Illumina HiSeq3000 platform (2 x 150 bp) were conducted at the Max-Planck-Genome-Center, Cologne, Germany (http://mpgc.mpipz.mpg.de).

### Genome assembly and Single Nucleotide Variants calling

PacBio HiFi reads from Zpa796 were assembled with IPA HiFi Genome Assembler v1.8.0, while Illumina reads for the 42 isolates of *Z. passerinii* and two technical replicates were assembled with SPAdes v3.11.1 (Bankevich *et al*., 2012). Transposable elements (TEs) were identified with REPET v3.0 (Quesneville *et al*., 2003, 2005; Flutre *et al*., 2011; Ahmed *et al*., 2011), and gene annotation was done with BRAKER (Hoff *et al*., 2019; Brůna *et al*., 2021).

SNV calling was performed with the GATK pipeline. After Identity By State (IBS) clone correction and data filtering, we retained 65 genotypes and 1,167,977 SNVs for five species of Zymoseptoria (Figure 2A). A detailed description of data processing is available in the Supplementary information.

### Population genomic analyses and interspecific relationships

The interspecific relationships were evaluated with SplitsTree (Huson & Bryant, 2006) with a set of pruned SNVs for non-clonal genotypes shared among the five species of the *Zymoseptoria* genus included in this work. Genetic relationships and population structure were assessed with a principal component analysis (Figure S6) with the R package SNPRelate (Zheng *et al*., 2012) and ADMIXTURE (Alexander & Novembre, 2009). Non-linked SNVs (r2 ≤ 0.25) were used for the ADMIXTURE analysis, spanning from K=1 to K=8, with ten replicates for each K value. Next, we computed nucleotide diversity and paired divergence index (*F_ST_*) between domesticated- and wild-host infecting populations with vcftools, with the flag-haploid in 10 Kb windows and steps of 2 Kb for *F_ST_*.

Protein sequences annotated with BRAKER for Illumina assemblies were used as input for the program Orthofinder v.2.5.5 (Emms & Kelly, 2019) for 21 isolates, 12 isolates of *Z. passerinii*, two for each *Z. ardabiliae*, *Z. pseudotritici* and *Z. tritici,* one for *Z. brevis* and two isolates for *C. beticola* which were used as outgroup. We built two phylogenies, the first with RaXML v8.2.9 (Stamatakis, 2014) using the parameters -m PROTGAMMAWAG -f a -x 321 -N 500 -p 123, and the second with IQ-TREE version 1.6.12 (Nguyen *et al*., 2015) with the flags --runs 10 -bb 1000 -nt 8 -m TEST. The best phylogenetic tree was visualized with FigTree v1.4.4.

### Virulence survey of *Z. passerinii* in *Hordeum*

We surveyed the virulence of *Z. passerinii* in the domesticated barley varieties Simba, Paradiesegerste and Golden Promise *(H. vulgare* ssp. *vulgare* henceforth *H. vulgare*); and the wild barley accessions: *H. jubatum* (seeds obtained from the Botanical Garden Halle, Germany); HOR2680 (*H. vulgare* ssp. *spontaneum,* Iran. Henceforth *H. spontaneum*) and GRA3223 (*H. murinum* ssp. *glaucum,* Armenia. Henceforth *H. murinum*), seeds were provided by the Leibniz-Institut für Pflanzengenetik und Kulturpflanzenforschung (IPK), Gatersleben, Germany. *Hordeum* plants were grown in a greenhouse in controlled conditions (∼20°C (day)/∼12°C (night) and a 16-hr day/8-hr night). Fungal strains were recovered on YMS agar and then transfer to YMS liquid media for four days at 18°C, plant inoculation was performed after the seedlings had fully developed the second leaf, following the methods described in (Fagundes *et al*., 2020). Samples for microscopy were gathered on days 4, 7, and during the transition from the biotrophic to the necrotrophic phase (day 9 for Golden Promise and day 10 for GRA3223) days post inoculation (dpi). Meanwhile, leaves for the quantitative virulence assay were collected at 21 dpi.

### Image analysis for quantification of virulence of *Z. passerinii* in *Hordeum*

### sp

Inoculated leaves from the three cultivar varieties of barley and the wild accessions HOR2680 and *H. jubatum* were collected at 21 dpi and mounted on paper sheets A4 with a reference mark and ID. High-resolution pictures (2400 dpi) were taken for each leaf. The cross virulence assay for Golden Promise (*H. v*ulgare spp. *vulgare*), HOR2680 (*H. spontaneum*), and GRA3223 (*H. murinum*) were analyzed with the frame and macro v2.1.1 from (Karisto *et al*., 2018). Leaves were scanned at 1,200 dpi with a Canon CanoScan LiDE 300. Disease severity was evaluated based on pycnidia/cm^2^ leaf surface and the percentage of leaf area covered by lesions (PLACL) (Figure 3C).

### Characterization of *Z. passerinii* infection in barley leaves by Confocal Laser Scanning Microscopy

Three leaves per time point (4, 7, shift day: 9 for *H. vulgare* and 10 for *H. murinum*, and 13 dpi) were harvested from each pathosystem (*H. vulgare*-Zpa63 and *H. murinum*-Zpa796) and mock-inoculated plants for the same time-points. Plant material was stained following a modified protocol from Haueisen, et al. (2019). Leaves were destained with 99% ethanol until cleared. Subsequently, a 1 cm piece of each sample was washed once with 1 mL of 1X phosphate-buffered saline (PBS, pH 7.4). Samples were incubated with 1 mL 10% KOH at 85°C to increase permeability and washed 3 times with 1 mL PBS to neutralize. The staining solution consisted of 0.02 % Tween-20 in 1X PBS, 10 µg/mL wheat germ agglutinin conjugated to fluorescein isothiocyanate (WGA-FITC), and 20 µg/mL propidium iodide (PI). Samples were vacuum infiltrated with 1 mL of staining solution for 10 min at 800 mbar followed by the slow release of the vacuum, and incubated overnight at 4°C protected from the light. The staining solution was replaced with 1 mL of 1X PBS and subjected to imaging or stored at 4°C.

An argon laser at 488 nm was used for excitation of FITC, and fluorescence was detected between 500 and 540 nm, while PI emission was detected from 600 to 670 nm when excited by a diode-pumped solid-state laser at 561 nm. Image stacks were obtained with an x/y scanning resolution of 1800 x 1800 pixels with a Zeiss Airyscan LSM 880. The visualization and processing of images were carried out utilizing and ZEN Black and Zen Blue software (Carl Zeiss Microscopy).

## Results

### Two new hosts of *Z. paserinii* in Northwest Iran

Previous reports have identified multiple hosts for *Z. passerinii,* including *H. vulgare, H. jubatum, H. nodosum,* and *H. distichon (Cunfer, 2000; Goodwin & Zismann, 2001; Seifbarghi et al., 2009; Quaedvlieg et al., 2011)*. For the leaf samples obtained from Iran, we were only able to identify the host at the genus level as being *Hordeum* spp. To further characterize the host of the *Z. passerinii* isolates, we conducted molecular identification with barcode markers. Our analysis revealed that *H. murinum* and *H. bulbosum* are also hosts for *Z. passerinii* (Table S2, Figure S1). These results expand the documented host range for *Z. passerinii* in Iran (Seifbarghi *et al*., 2009). Based on this finding, we addressed whether the samples collected from different hosts comprise a unified genetic pool or have evolved into host-specific lineages.

### De novo assembly of Zpa796 reveals accessory chromosomes in wild-host infecting *Z. passerinii*

Before this study, only one long-read assembly of *Z. passerinii* from domesticated barley had been sequenced and assembled (Feurtey *et al*., 2020). Previous comparative genome analyses of multiple isolates of the sister species *Z. tritici* have revealed the presence of multiple accessory chromosomes (Goodwin *et al*., 2011; Möller *et al*., 2018). To survey the presence of accessory chromosomes in *Z. passerinii*, we generated a reference assembly for *Z. passerinii* based on the long-read sequencing of the Iranian isolate Zpa796. The resulting PacBio assembly comprise a total lenght of 31.38 Mb, including 31 contigs; 12 of them with telomeres at one extreme, while contigs 3 and 17 were assembled from telomere to telomere (Figure S2). A Benchmarking Universal Single-Copy Orthologs (BUSCO) v5.3.2 analysis revealed ∼ 98 % of genome completeness using the lineages fungi and ascomycota (Table 1). Meanwhile, the transcript prediction based on RNA performed with BRAKER identified 11,727 gene models and 10,461 proteins, of which 977 were predicted to be secreted by interproscan. Repeat sequences coverage in the Zpa796 genome, as predicted with REPET, was 13.3 % (Table 1).

**Table 1.** Metrics of Zpa796 genome assembly.

| Feature | Value |
| --- | --- |
| Isolate | Zpa796 |
| Origin | East Azarbaijan, Iran |
| Host | Hordeum murinum |
| Year of field collection and isolation | 2018, 2020 |
| Contigs number | 31 |
| Total Length (bp) | 31,389,365 |
| N50 | 1594512 |
| L50 | 8 |
| Contigs with telomeric repeats on both ends | 2 |
| Complete BUSCO genes (%) | 98.7 |
| Number of gene models | 11,702 |
| TE coverage (%) | 13.3 |
| Mean contigs size (Mb) | 1.012 (SD = 0.87) |

We compared full genome PacBio assemblies, from the wild-grass isolate Zpa796 with the already available reference genome of Zpa63 from domesticated barley, Zpa63 (Feurtey *et al*., 2020). The genome alignment revealed that ten contigs, ranging in size from 0.1 to 1.4 Mb, did not align with the Zpa63 genome; four of them have telomeres at one extreme (Figure S2). The occurrence of these long chromosome fragments present in Zpa796, but not Zpa63 suggests the presence of accessory chromosomes also in *Z. passerinii* (Fig 1A).

**Figure 1.**
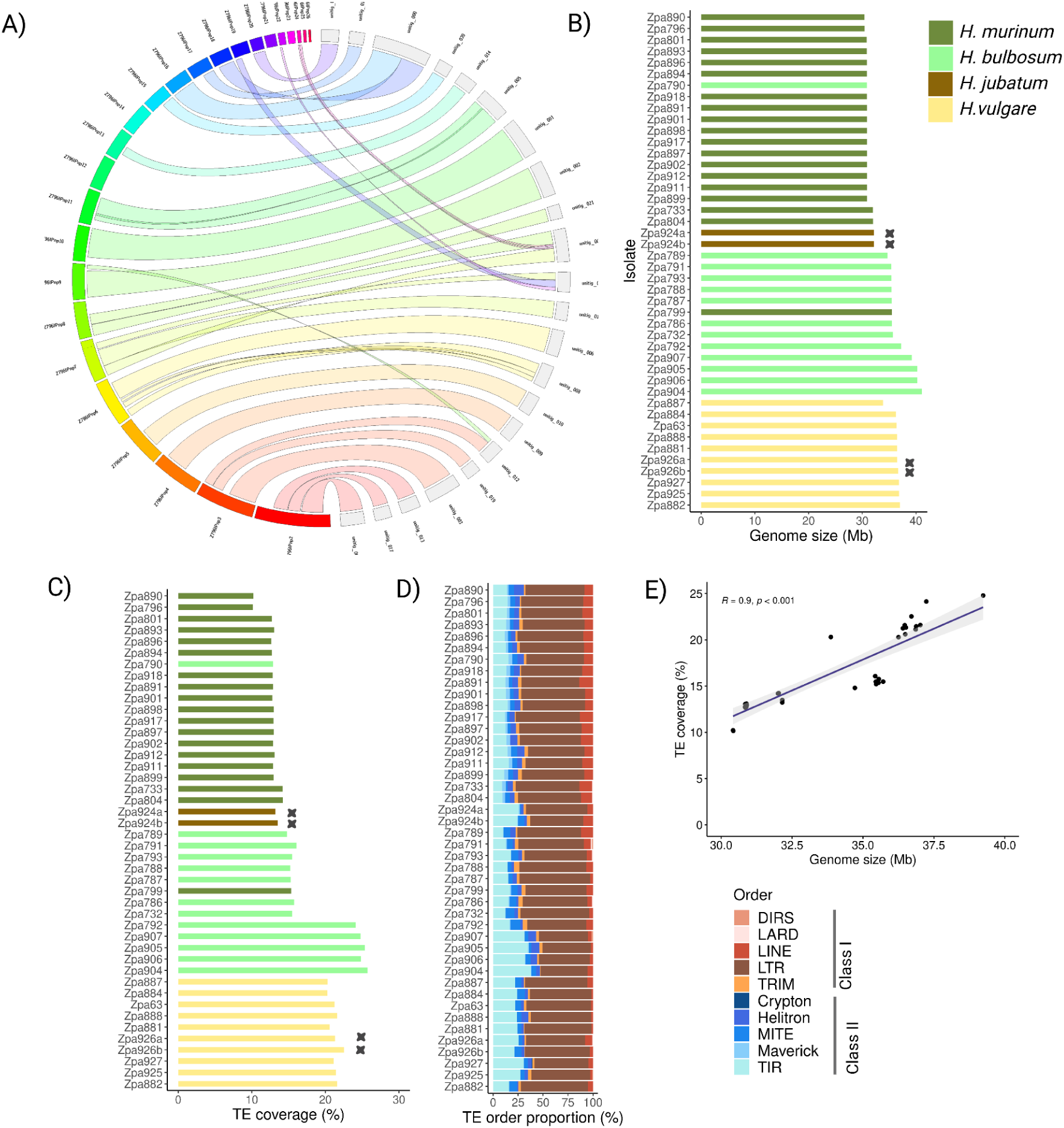
Intraspecific genomic variation of *Z. passerinii*. Panel A) Comparison of PacBio genome assemblies between Zpa796 (on the right) and Zpa63 (on the left); B) Genome assembly length of the *Z. passerinii* isolates; C) Percentage of TEs coverage per genome assembly, crosses on the bar plot indicate technical replicates and D) Proportion of TEs orders.

### Accessory segments of wild and domesticated-hosts infecting populations are enriched in TEs

Due to the presence of an accessory genome and the genome size difference between Zpa63 and Zpa796, we decided to survey genome size variation among *Z. passerinii* populations. We built de novo Illumina genome assemblies for the 42 isolates, non-clone corrected, of *Z. passerinii*. Furthermore, we performed a detailed analysis of genetic variation among and within populations of *Z. passerinii* based on SNVs identification.

The Illumina assembly length of the reference isolate Zpa796, 30.42 Mb, corresponded to the genome length that we obtained from the de novo assembly with PacBio (31.38Mb). Interestingly, we observed a genome size variation among isolates from the same and different hosts (Fig 1B). The isolates Zpa796 and Zpa890 were obtained from the same locality but different plants and exhibited the smallest genomes (∼ 30 Mb) (Figure 1B). Interestingly, we did not observe a correlation between genome size and the host species. In fact, the largest genome size variation (32–41 Mb) was between isolates obtained from a single host, *H. bulbosum* indicating a large plasticity in genome content in *Z. passerinii* from both wild and domesticated hosts. The genome size of each isolate was positively correlated with the predicted TE content (Figure 1E), underlining the relevance of TEs in driving genome size evolution in *Z. passerinii*.

TEs were annotated, and a comparative analysis was performed on sequences assigned to a class (63.02 %); unclassified (32.27 %) and potential host genes (4.7 %) sequences were omitted. (Figure S3). TEs from Class I were the most abundant for all the genomes, with a mean proportion of 70.84 % versus 29.16 % for Class II (Figure 1D). LINE (class I) and Maverick orders (class II) were statistically significantly more abundant in wild-host infecting isolates (Wilcoxon rank test, p-value < 0.01). The four isolates with the longest genome, Zpa 904 to Zpa 907 (39–41 Mb), harbored a higher proportion of TIR order (31–37 %) (Figure 1D, <u>Table S</u>5).

### Domestication of the host has led to divergence among *Z. passerinii* populations

The wheat pathogenic fungus *Z. tritici* emerged in the Middle East and subsequently evolved and spread with its host, a scenario referred to as host-tracking (Stukenbrock et al. 2011). To investigate if barley-infecting lineages of *Z. passerinii* also arose through host tracking, we explored genetic variation and divergence at the population level. First, we evaluated the interspecific relationships to verify that the isolates of *Z. passerinii* group into a single species. To achieve this, we first performed genotype-filtering and clone-correction based on a collection of Illumina sequenced *Zymoseptoria* genomes, including the *Z. passerinii* genomes generated here (see Materials and Methods). As a result, we kept 65 out of 93 genotypes for the five species of *Zymoseptoria* included in our analyses, and a total of 1,167,977 SNVs. The interspecific relationships were surveyed with a phylogenetic network analysis (Figure 2A). Independent of sampling location and the original host, all the isolates from *Z. passerinii* clustered in a single node of the network (Figure 2A). The only exception to this was the isolate Zpa924 collected in Minnesota, USA, from *H. jubatum* (Table S3); which was previously reported as to be *Z. passerinii* (ID P26515) (Goodwin & Zismann, 2001). However, based on our whole-genome analyses, we found that Zpa924 very likely belongs to a hitherto unknown *Zymoseptoria* sister species, as it does not cluster with any of the other four species included in this work. Due to its outlier behavior, Zpa924a and its technical replicate (Zpa924b) were removed from the downstream population analyses.

**Figure 2.**
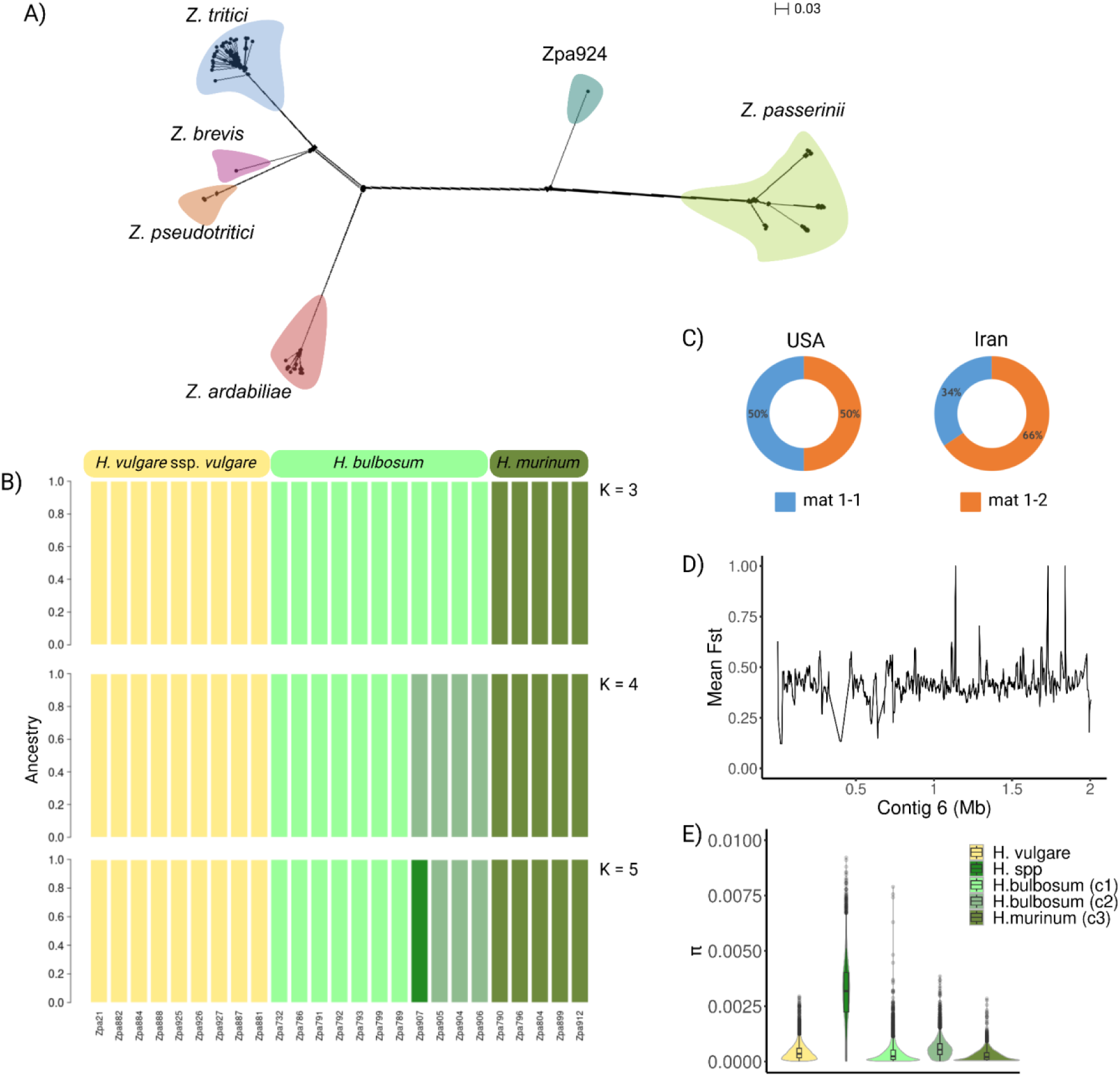
Population genomics for *Z. passerinii* isolated from domesticated barley (and wild barley. A) Phylogenetic network based on SNVs shared among five *Zymoseptoria* genus species; B) Admixture analysis for the 25 non-clonal *Z. passerinii* genotypes from domesticated (yellow) and wild *Hordeum* (green tones), K= 3 had the lowest cross-validation error; C) Mating types proportion for Z. passerinii isolates sampled in the USA and Iran; D) Divergence index (*F_ST_*) along contig 1 between domesticated- and wild-host infecting genotypes; and E) The first two violin plots show nucleotide diversity for domesticated- and wild-host infecting genotypes, the last three box plots show π, for the three genetic clusters (c1, c2, c3) identified for isolates from wild *Hordeum*, the colors match the corresponding ancestry.

After verifying the species clustering, an independent filtering was performed for the population genomic analyses. When excluding the sister species and applying the same filters, we retained 2,205,972 SNVs only for *Z. passerinii*. Interestingly, despite the manual clonal correction with SSR markers performed for *Z. passerinii* isolates, we observed high clonality among genotypes from the same leaf (<u>Fig, S</u>4). To avoid confounding effects for the population genetic analyses, we conducted further analyses with an even more reduced dataset. We kept 16 out of the 33 isolates from the wild *Hordeum* hosts, including 11 isolates from *H. bulbosum*, and 5 from *H. murinum*. Additionally, we kept the nine isolates from domesticated barley; these were obtained from plant pathogen collections and therefore represents a broader temporal and spatial sampling scale.

The clustering pattern of the phylogenetic network was corroborated by the Admixture analysis. We observe a deep population structure with no signs of admixture, where K = 4 had the lowest cross-validation error (Figure S7). Despite the isolates obtained from *H. vulgare*, were collected in different years and at different counties in the USA (Figure S10); we detected a single cluster when plotting from K = 3 to K = 5. When K = 4, we observe three clusters for the wild-host infecting isolates: for *H. bulbosum* isolates, we detected two ancestries that matched the locations, Marand and Arasbaran Forests counties in Iran, and sampling years, 2018 and 2020, respectively. While we observed a single cluster for *H. murinum*, sampled in 2018 at two locations (Table S3, Fig 2B). These findings indicate a strong effect of host specificity in population subdivision and suggest that little gene flow occurs between *Z. passerinii* populations adapted to distinct *Hordeum* hosts.

Based on the type of host, domesticated and wild barley, we computed population divergence (*F_ST_*) and nucleotide diversity (π). The mean *F_ST_* across the genome was ∼ 0.4 (Figure 2D and Figure S9), which suggests that barley domestication and host specialization may have been drivers of population divergence. The levels of genetic diversity were significantly higher for the wild-host *Z. passerinii* populations, however when π was computed per genetic cluster, the π values were lower and closer to the π value of the domesticated-host *Z. passerinii* population (Fig 2E).

New genetic combinations can be generated by sexual reproduction, therefore we addressed the putative contribution of sexual mating in the *Z. passerinii* populations. To this end, we surveyed the distribution of mating type alleles for all the isolates. *Zymoseptoria passerinii* is a heterothallic fungus and mating occurs between isolates of opposite mating types, mat 1-1 and mat 1-2 alleles (Goodwin *et al*., 2003). We surveyed the mating type alleles distribution and found that mat 1-1 and mat 1-2 have a frequency of 50% each in the USA, while mat 1-1 was present at 34 % vs 66 % for mat 1-2 of the Iranian isolates (Fig 2C). Even though mat 1-2 was more abundant than mat 1-1, both mating types were found in all sampling localities, except in Ardabil-Khalkhal county, Iran, from which we have only two clonal isolates with the same mating type. Interestingly, the genotypes Zpa799 and Zpa801, both from the same location (Marand-Tabriz road) and isolated from the same lesion (IEAMSS1_1) were classified as clonal based on the IBS filter, nonetheless the blast analysis showed that they have different mating types (Table S3). The last implies that despite a high degree of inbreeding, which would result in high IBS (> 0.999) different mating types are present in natural populations of *Z. passerinii* and sexual reproduction contributes to the population genetic structure of the pathogen populations.

### Host specific-lineages of *Z. passerinii* in natural ecosystems

The population structure associated with the host species (Figure 2B) and the high divergence index (*F_ST_* ∼ 0.4) of *Z. passerinii* from wild and domesticated hosts (Figure 2D); suggested host specialization or incipient speciation process in *Z. passerinii*. We expeceted the geographic distance between the populations of *Z. passerinii* obtained from domesticated and wild hosts to contribute to genetic divergence. However, the collection from Iran reveal that other factors are also at stake. The wild isolates were collected in sympatry in Northwest Iran but on different hosts. Host specialization occurs when a pathogen shifts to a new host and undergoes an adaptation process, resulting in subpopulations better fitted to the new host. This can ultimately change the ability to infect the original host (Thines, 2019). In order to further evaluate the evolution of host-specific lineages, we used both phylogenetic and experimental approaches. First, we built a phylogeny using 6,563 single copy of orthologs identified for the five species of *Zymoseptoria* genus included in this work, and the pathogen *C. beticola* here used as an outgroup. Interestingly, *Z. passerinii* isolates clustered with a support of 100 on three distinct branches corresponding to the three host species from which they were isolated. The same phylogenetic arrangement and support were obtained with the two methods using RAxML and iqtree, providing further evidence of host-specific lineages of *Z. passerinii*. Moreover, as suggested by the phylogenetic network, the isolate Zpa924 was placed in a separate branch that is closer to *Z. passerinii* in comparison to the other four *Zymoseptoria* species, providing further support to our hypothesis that Zpa924 most likely represents a sister species of *Z. paserinii* (Fig 3A).

**Figure 3.**
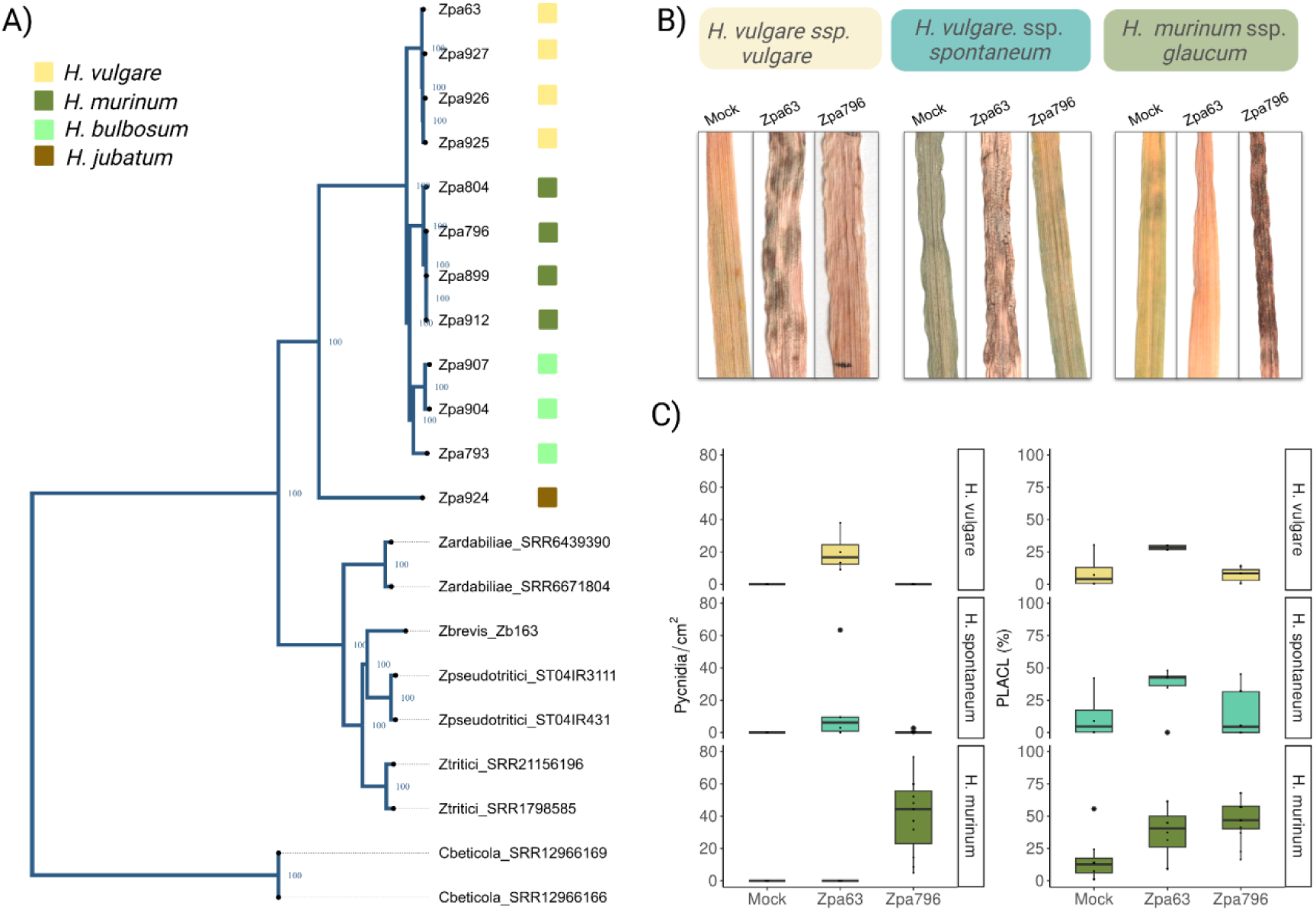
Evidence of host-specific lineages. A) Phylogenetic tree for *Zymoseptoria* species rooted with *Cercospora beticola* and based on 6,563 single copy orthologous genes. The values show the bootstrap support for each branch, and the colored squares show the host from which each genotype of *Z. passerinii* was isolated. B) In planta phenotypic assay of *Z. passerinii* in domesticated barley (*H. vulgare* ssp. *vulgare*, cultivar Golden Promise), the barley wild ancestor (*H. sponateum*, HOR2680) and the CWR *H. muruninum* (GRA3223). C) Virulence quantitative assay of *Z. passerinii* in domesticated *Hordeum* (cultivar Golden Promise), and the CWRs *H. spontaneum* (HOR2680) and *H. murinum* (GRA3223) with the features pycnidia/cm2 of leaf and Percentage of Leaf Area Cover by Lesions (PLACL).

To further test the extent of host specialization experimentally, we surveyed the host range of *Z. passerinii* using ten isolates; six from the domesticated host (*H. vulgare*) and four from the three wild hosts (*H. murinum*, *H. bulbosum* and *H. jubatum*). We note that wild-host infecting isolates tend to grow primarily as hyphae and melanize before reaching high-density cultures. Thus, we selected wild-host infecting isolates according to their ability to grow as single cells in suspension without forming melanized aggregates in YMS liquid media, which otherwise would have prevented the use of consistent inocula for the different isolates. We assessed virulence using i) pycnidia numbers per cm2 of leaf and ii) PLACL on inoculated leafs. The six domesticated-host infecting isolates: Zpa63, Zpa881, Zpa882, Zpa884, Zpa887, and Zpa888 were able to infect and produce pycnidia in the three cultivated varieties tested of *H. vulgare*: Golden Promise, Simba and Paradiesegerste, and four isolates produced pycnidia in the Iranian accession of *H. spontaneum* (HOR2680) (<u>Table</u> <u>S</u>6). In contrast, none of the four isolates from wild *Hordeum* species —Zpa789, Zpa799, Zpa796, and Zpa924— were able to produce pycnidia on the tested cultivars, or *H. jubatum*, even when Zpa924 was isolated from this wild *Hordeum* species (Figure 3A).

On an independent cross virulence assay with Zpa63 (domesticated-host) and Zpa796 (wild-host), we used Golden Promise, *H. spontaneum* (HOR2680, Iran) and *H. murinum* (GRA3223) seeds from Armenia, which is geographically close to Northwest Iran, where this pathogen naturally occurs. Zpa63 produced significantly higher pycnidia density and necrosis on Golden Promise (*H. vulgare*) and HOR2680 (*H. spontaneum*), while Zpa796 did not produce any symptoms on *H. vulgare* and *H. spontaneum*. In contrast, Zpa796 produced a high pycnidia density on the GRA3223 accession (*H. murinum*). Manual leaf evaluations revealed no pycnidia in the controls or in leaves of *H. murinum* innoculated with the isolate Zpa63. Zpa63 induced necrosis in all three hosts (compared to mock samples) although not sporulating. The isolate Zpa796, on the other hand, did not induce necrosis *H. vulgare* but induced some necrosis in *H. murinum* (**Fig 3C**). The incapability of isolates originating from wild barley to infect the *H. vulgare* cultivars supports a scenario of host specialization driving the genetic divergence of *Z. passerinii* populations.

### Persistence of *Z. passerinii* in natural ecosystems is linked to environmental conditions

The last outbreak of *Z. passerinii* in a domesticated ecosystem was reported in 2006 (Toubia-Rahme & Steffenson, 2004), although our sampling demonstrates that the pathogen persists in natural ecosystems. Eve more, under control conditions, the domesticated isolates were able to infect current cultivars of barley (Table S6). Therefore, we speculate whether the persistence of *Z. passerinii* in natural ecosystems is linked to a different disease onset observed in wild-host infecting lineages compared to the domesticated-host. To evaluate this, we surveyed with CLSM the infection development in the domesticated pathosystem, Zpa63 in *H. vulgare* (Golden Promise), and the wild pathosystem, Zpa796 in *H. murinum* (GRA3223).

Our microscopy analyses show that the SSLB disease progression and timing are highly correlated and comparable to the hemibiotrophic infection development described for *Z. tritici* in *Triticum aestivum* (Haueisen, et al. 2019). After foliar inoculation, we observed that both strains established a biotrophic infection characterized by hyphae’s symptomless colonization of the mesophyll layers (Figure 4 A, B). After 9 or 10 dpi, there was a shift towards a necrotrophic phase characterized by leaf necrosis and the development of pycnidia (Figure 3B, C) and the accumulation of hyphae in the substomatal cavities (Figure 4C) that subseqently give rise to conidiogenous cells (Haueisen, et al. 2019). Finally, mature pycnidia developed in the substomatal cavities surrounded by hyphae and collapsed mesophyll cells (Figure 4D). The similarities between the two pathosystems support that the persistence of SSLB in natural ecosystems is not due to a different disease development.

**Figure 4.**
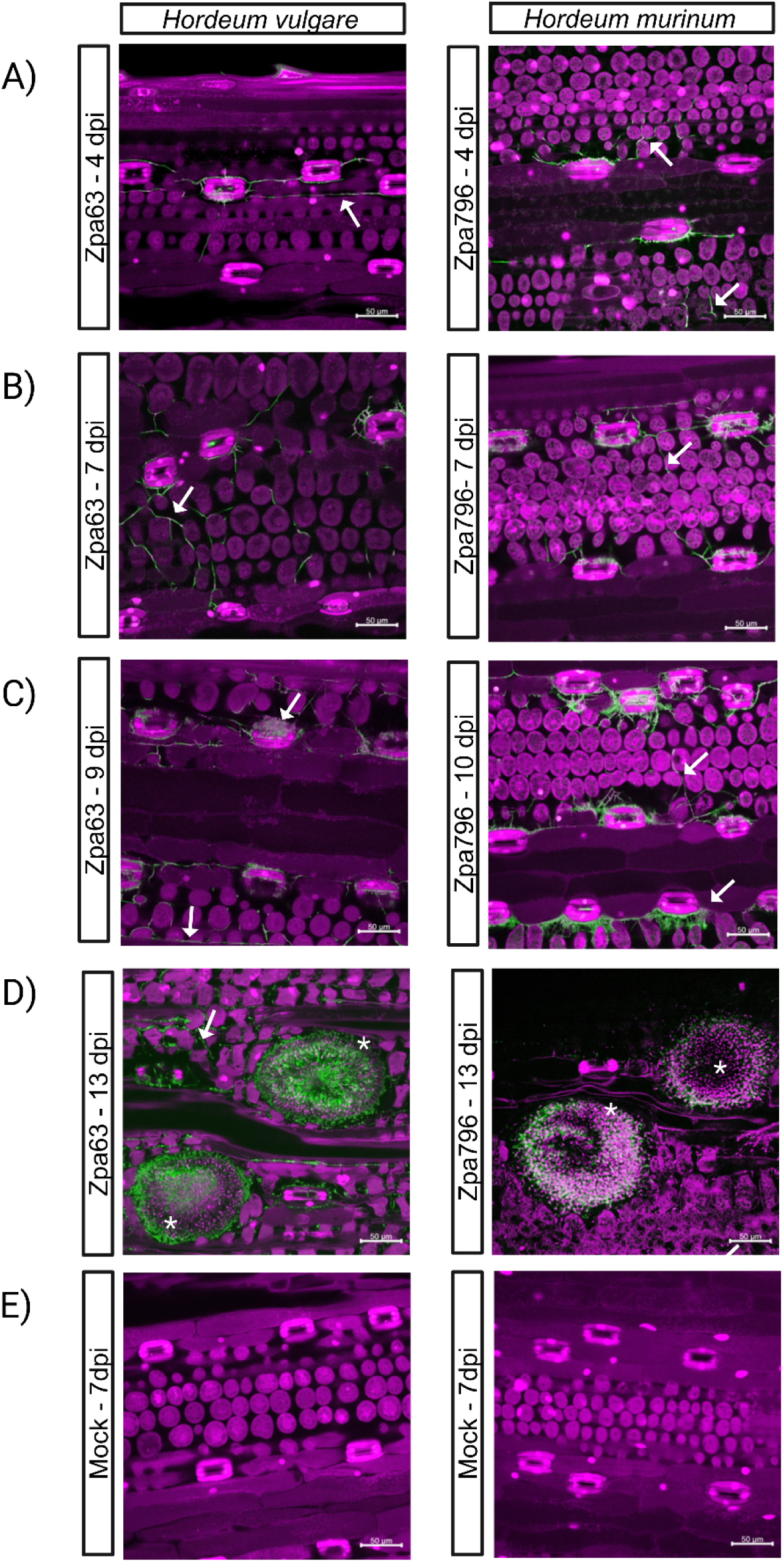
Comparison of infection development of *Z. passerinii* in *H. vulgare* (domesticated host) and *H. murinum* (wild host). The fluorescence confocal micrographs show the infection of Zpa63 and Zpa796 in their specific host at 4 dpi (A), 7dpi (B), 9 or 10 dpi (C), 13 dpi (D), and the Mock inoculation at 7 dpi for comparison (E). Plant cells and nuclei appear in purple while hyphae appear in green (white arrows), white asterisks indicate mature pycnidia. The scale bar represents 50 um.

## Discussion

This study examines the evolutionary dynamics of *Z. passerinii*, a sister species of the plant pathogen *Z. tritici*, in a natural ecosystem in Northwest Iran. We documented SSLB disease persistence and population genetic processes driving the emergence of host-specific lineages in wild barley. SSLB disease, as studied under control experimental conditions, exhibits significant similarities in domesticated and wild hosts. We speculate that development of SSLB is mainly dependent on favorable environmental conditions. Given the close relatedness of *Z. passerinii* lineages infecting different *Hordeum* species, a limited set of genes may regulate host specialization. Additionally, disease timing and regulation are likely conserved across different hosts. Considering the significant economic importance of *Z. tritici* in wheat, we propose the *Z. passerinii*-barley pathosystem as a valuable tool for studying septoriosis, as it simplifies the complexities associated with host genetics.

### Did barley-infecting *Z. passerinii* lineages emerge through host tracking or host range expansion?

The phylogenetic analyses which we performed based on 6,563 single copy of orthologs support that domesticated and wild-host infecting lineages of *Z. passerinii* comprise a single species. However, the intraspecific divergence among genetic clusters of *Z. passerinii* in the USA and Northwest Iran (Figure 2B,D) contrasts with the shallow population structure described for the sister species *Z. tritici,* for which divergence is observed mainly at the continental level (Zhan *et al*., 2003; Feurtey *et al*., 2023)). Furthermore, we observed that domesticated-host infecting isolates of *Z. passerinii* were able to infect and produce SSLB disease in *H. vulgare* and *H. spontaneum* (HOR2680, Iran), the closest wild ancestor of barley. This observation supports the hypothesis that CWRs can serve as reservoir for crop plant pathogens (Papaïx *et al*., 2015). More importantly, this finding suggests that barley domestication did not lead to the emergence of a host specialized lineage, as it has been documented for the sister species *Z. tritici* in wheat, but rather through an expansion of the host range of a lineage probably already able to infect *H. spontaneum*.

The fact that the American isolate Zpa924 could not infect its natural host species, *H. jubatum* (Goodwin & Zismann, 2001), suggests that pathogenicity is a product of local adaptation to specific host populations. This assumption gains support because Zpa796 successfully induced SSLB disease in *H. murinum* ssp. *glaucum* (GRA3223), an accession collected in Armenia close to the natural pathogen population, highlighting a putative role of local adaptation in host-pathogen interactions.

### Intraspecific genomic variation of *Z. passerinii*

The genome assembly of Zpa796 resulted in a genome smaller than that of Zpa63. Intriguingly, ten contigs did not align with Zpa63, reinforcing the presence of accessory chromosomes in *Z. passerinii*, a trait to be conserved across the *Zymoseptoria* genus (Stukenbrock *et al*., 2011). It has been speculated that accessory genomes shared across the *Zymoseptoria* genus likely serve as sources of structural and genetic variation through mitotic division (Möller *et al*., 2018), evolving at a faster pace than the core genome.

On the other hand, the correlation between genome length and TEs coverage in *Z. passerinii* isolates contrasts to *Z. tritici,* where an increase in TEs content correlated with continental spread of the wheat pathogen (Oggenfuss *et al*., 2021; Lorrain *et al*., 2021). In *Z. passerinii* however, TEs content varies across both continental and local scales. For instance, populations infecting wild hosts in Northwest Iran, located at distances of 14 to 222 Km, exhibit substantial variation in genome length and TEs content. Notably, Class I TEs (retrotransposons) make up 70.84% of the repetitive sequences in the genome of *Z. passerinii*, mirroring findings for *Z. tritici (Dhillon et al., 2014)*. This aligns with earlier evidence demonstrating that the accessory genome of *Z. tritici* is enriched in repetitive elements and noncoding sequences (Dhillon *et al*., 2014). Additionally, based on observations for other Dothideomycetes species (de Wit *et al*., 2012), retrotransposons likely have contributed to the genome expansion of *Z. passerinii* and this event is happening at small geographic scale among populations located in a linear distance of hundreds of kilometers.

### Wild-host infecting populations shed light on species delineations

Despite previous reports positioning *Z. pseudotritici* as a sister species to *Z. trititici (Quaedvlieg et al., 2011; Stukenbrock et al., 2012b),* our phylogenetic analysis, employing both SNVs networks and single-copy ortholog trees, reveals a closer relationship between *Z. pseudotritici* and *Z. brevis* than with *Z. tritici.* Our study represents the most comprehensive phylogenetic analysis within the genus, utilizing, 6,563 single-copy orthologs.

While *Z. tritici* and *Z. pseudotritici* have traditionally been considered sister species (Stukenbrock *et al*., 2012a,b), the separation among *Z. tritici*, Z*. pseudotritici*, and *Z. brevis* has lacked conclusive support (Bayesian posterior probabilities = 0.75) (Stukenbrock *et al*., 2012b). Contrarily, our analysis obtained full support (100%) for the clade distribution, highlighting *Z. brevis* and *Z. pseudotritici* as sister species. Moreover, including new genotypes of *Z. passerinii* isolated from wild-hosts revealed that Zpa924, previously classified as *Z. passerinii*, represents a distinct species from *Z. passerinii*; most likely *Z. halophila*, a sister species to *Z. passerinii* (Quaedvlieg *et al*., 2011; Stukenbrock *et al*., 2012b). Nonetheless, extensively characterization of phenotypic traits is required to consolidate this finding.

### Does host-specificity in sympatric populations lead to population divergence in natural ecosystems?

The wild-host infecting populations exhibit distinct ancestries in geographic sympatry, geographic range of 14-222 km (Figure S10), which were related to host species (Figure 2B). Conversely, the domesticated-host infecting isolates, comprising genotypes from four counties in North Dakota and Minnesota, formed a separate cluster clearly distinct from wild-host infecting isolates, suggesting an absence of gene flow at local and continental scales. This population structure may be driven by different processes; we speculate that geographic isolation and the distribution of resistant barley cultivars likely shaped the divergence of domesticated-host infecting populations, while reproductive isolation and host specialization has influenced wild-host infecting populations in Northwest Iran.

Population genetic analyses have revealed that high within-field genetic variation observed in the sister species *Z. tritici* results from recurrent sexual reproduction cycles and continuous gene flow (Zhan *et al*., 1998; McDonald *et al*., 2022). In contrast, *Z. passerinii* exhibits low genetic diversity within populations and high divergence among genetic clusters (Figure 2D, E). This supports the notion of local adaptation occurring among populations in sympatry, likely driven by the availability of multiple hosts in Northwest Iran. Interestingly, the diversity of genotypes per lesion observed for *Z. passerinii*, resembles that of *Z. tritici (Linde et al., 2002)*. We isolated 16 non-clonal genotypes from five lesions, meaning an average of 3.2 different genotypes per lesion.

Additionally, we observed an even distribution of mating type alleles across the sampled regions in the USA (Figure 2C), which agrees with previous results for *Z. passerinii* in the northern USA (Lee & Neate, 2007a). A dissimilar ratio of mating type alleles in Northwest Iran suggests that sexual reproduction and migration of genotypes is less frequent than in the agricultural environment, further supported by the linkage disequilibrium pattern of wild-host infecting populations (Figure S5). Nonetheless, we acknowledge the high clonality degree and sampling size of populations can not be discarded as a cause of the LD pattern observed. Moreover, we note that the Northwest Iranian isolates were found in sympatry. Therefore, the infrequency of sexual reproduction and the strong divergence between host-specific lineages could also be a consequence of reproductive isolation. However, this scenario should be properly tested through controlled crosses.

### Does SSLB persist in natural ecosystems due to differing disease onset in wild versus domestic hosts?

SSLB disease in domesticated barley is considered to be dependent on warm climates (20 to 24°C) and extended periods of humidity (St Pierre *et al*., 2010); when the spontaneous outbreaks of SSLB occur these can cause yield losses from 20 to 40 % and reduce the suitability of barley for malting (Green & Bendelow, 1961; Van Ginkel *et al*., 1999). The sporadic nature of SSLB contrasts with *Z. passerinii* virulence under control conditions for domesticated and wild hosts. Despite our speculation that a different onset and timing of disease in wild hosts could be responsible for the persistence of this pathogen in natural ecosystems; the comparison of the domestic and wild pathosystems, in greenhouse conditions, showed a similar development of the disease, which suggest environmental conditions have a main role as described in the disease triangle: pathogen, host and environment (McNew, 1960).

It has been suggested that in ascomycetes, adaptation to host and habitat can lead to reproductive isolation (Giraud & Gourbière, 2012; Stukenbrock, 2013). This was indeedn documented for *Z. tritici* for which high differentiation in sympatry associated with host specialization led to the emergence of a new pathogen *(Stukenbrock et al., 2007)*. This scenario resembles domesticated host infecting *Z. passerinii* isolates, which can infect the wild ancestor of barley but fail to infect more distantly related species like *H. murinum* ssp. *glaucum* and *H. jubatum*. The domestication of *H. vulgare* resulted in its divergence from *H. spontaneum* approximately 8,000 ya (Badr *et al*., 2000; Morrell & Clegg, 2007), but despite this divergence, our results showed that infection mechanisms are shared for domesticated and wild *Hordeum*. However, the fact that domesticated-host isolates were unable to infect *H. murinum ssp. glaucum*, a more distant relative, that diverged from *H. vulgare* around 10 million ya (Jakob & Blattner, 2006) indicates the boundaries of host specificity for *Z. passerinii*. Thus, the observed divergence and the phylogenetic analysis of *Z. passerinii* support the emergence of sympatric host-specific lineages of the hemibiotrophic fungi *Z. passerinii* in wild hosts and a natural ecosystem.

## Acknowledgments

We thank the Leibniz-Institute für Pflanzengenetik und Kulturpflanzenforschung for providing us with wild barley accessions. We thank Prof. Dr. Gert Kema and Els Verstappen from Wageningen University, and Prof. Dr. Bruce McDonald and Julien Alassimone from ETH for providing us with isolates of *Z. passerinii*. Likewise, we thank Janine Müller, Susanne Braun, Doreen Landermann, Alina Waldmann, and Blanca Stuelpnagel for technical support for DNA extraction and clonal correction. We thank Frauke Caliebe for assistance in the virulence assays of the domesticated pathosystem. We thank Derk Wachsmuth and Kristian Ulrich for the technical support for performing the bioinformatic analysis on HPC of the MPI for Evolutionary Biology. Finally, we thank Dr. Danilo Pereira and Dr. Carolina Sardinha for the methodological and results discussion during the development of this research.

## Author contributions

I.C.R.B and E.H.S. Conceptualized the study

I.C.R.B Analyzed the genomic data, designed and performed virulence experiments, and wrote the paper.

V.F.N. Performed virulence experiments, the CML analysis and critical review the paper.

J.H. Performed *Z. paserinii* isolation from 2018

A.A. & F.S. Performed field sampling and phenotyping identification of *Hordeum* species

E.H.S. Funding acquisition, resources, and critical review All authors read, commented and approved the manuscript

## Data availability

Data will be made available for reviewers upon request and otherwise for publication of the results described in the manuscript.

## Supplementary Information

### Tables

Table S1. Primer sequences for clonal correction and host identification Table S2. *Hordeum* species identification

Table S3. Metadata of *Z. passerinii* isolates

Table S4. Public genomes used to perform the SNVs calling

Table S5. TEs proportion comparison for domesticated- and wild-host infecting isolates

Table S6. In planta inoculation experiments performed for Z. passerinii in four species of Hordeum.

### Figures

Figure S1. Maximum likelihood tree for *Hordeum* herbarium samples Figure S2. Contig size for the genome assembly of the isolate Zpa796

Figure S3. Proportion of repeat sequences annotated for 42 non-clonal corrected isolates of *Zymoseptoria passerinii*.

Figure S4. Identity By State (IBS) for A) *Z. passserinii*, B) *Z. tritici*, and C) *Z. ardabiliae*

Figure S5. Linkage Disequilibrium for the contig two of *Zymoseptoria passerinii*. Figure S6. Principal Component Analysis plot for *Zymoseptoria passerinii*

Figure S7. Mean cross-validation error (CVE) for ten admixture runs, bars represent the SD for each K-value.

Figure S8. Ancestry analysis for the 44 genotypes (clonal and non-clonal) of *Z. passerinii*, where the K = 6 has the lowest CVE.

Figure S9. Whole-genome divergence (*F_ST_*) for non-linked SNPS (r^2^ < 0.25) between *Z. passerinii* isolates from domesticated and wild hosts.

Figure S10 Geographical location of the 42 genotypes of Z. passerinii included in this study.

## References

Ahmed I, Sarazin A, Bowler C, Colot V, Quesneville H. 2011. Genome-wide evidence for local DNA methylation spreading from small RNA-targeted sequences in Arabidopsis. Nucleic acids research 39: 6919–6931.

Alexander DH, Novembre J. 2009. Fast Model-Based Estimation of Ancestry in Unrelated Individuals. : 1655–1664.

Allen GC, Flores-Vergara MA, Krasynanski S, Kumar S, Thompson WF. 2006. A modified protocol for rapid DNA isolation from plant tissues using cetyltrimethylammonium bromide. Nature protocols 1: 2320–2325.

Badr A, Müller K, Schäfer-Pregl R, El Rabey H, Effgen S, Ibrahim HH, Pozzi C, Rohde W, Salamini F. 2000. On the origin and domestication history of Barley (Hordeum vulgare). Molecular biology and evolution 17: 499–510.

Baldwin BG. 1992. Phylogenetic utility of the internal transcribed spacers of nuclear ribosomal DNA in plants: an example from the compositae. Molecular phylogenetics and evolution 1: 3–16.

Baldwin BG, Sanderson MJ, Porter JM, Wojciechowski MF, Campbell CS, Donoghue MJ. 1995. The its region of nuclear ribosomal DNA: A valuable source of evidence on angiosperm phylogeny. Annals of the Missouri Botanical Garden. Missouri Botanical Garden 82: 247.

Banke S, Peschon A, McDonald BA. 2004. Phylogenetic analysis of globally distributed Mycosphaerella graminicola populations based on three DNA sequence loci. Fungal genetics and biology: FG & B 41: 226–238.

Bankevich A, Nurk S, Antipov D, Gurevich AA, Dvorkin M, Kulikov AS, Lesin VM, Nikolenko SI, Pham S, Prjibelski AD, et al. 2012. SPAdes: a new genome assembly algorithm and its applications to single-cell sequencing. Journal of computational biology: a journal of computational molecular cell biology 19: 455–477.

Bieniek W, Mizianty M, Szklarczyk M. 2015. Sequence variation at the three chloroplast loci (matK, rbcL, trnH-psbA) in the Triticeae tribe (Poaceae): comments on the relationships and utility in DNA barcoding of selected species. Plant systematics and evolution = Entwicklungsgeschichte und Systematik der Pflanzen 301: 1275–1286.

Brůna T, Hoff KJ, Lomsadze A, Stanke M, Borodovsky M. 2021. BRAKER2: automatic eukaryotic genome annotation with GeneMark-EP+ and AUGUSTUS supported by a protein database. NAR genomics and bioinformatics 3: lqaa108.

Choi Y-J, Thines M, Runge F, Hong S-B, Telle S, Shin H-D. 2011. Evidence for high degrees of specialisation, evolutionary diversity, and morphological distinctiveness in the genus Bremia. Fungal biology 115: 102–111.

Cunfer BM. 2000. Stagonospora and Septoria diseases of barley, oat, and rye. Canadian journal of plant pathology. Revue Canadienne de phytopathologie 22: 332–348.

Dhillon B, Gill N, Hamelin RC, Goodwin SB. 2014. The landscape of transposable elements in the finished genome of the fungal wheat pathogen Mycosphaerella graminicola. BMC genomics 15: 1–17.

Eck JL, Kytöviita M-M, Laine A-L. 2022. Arbuscular mycorrhizal fungi influence host infection during epidemics in a wild plant pathosystem. The New phytologist 236: 1922–1935.

Emms DM, Kelly S. 2019. OrthoFinder: phylogenetic orthology inference for comparative genomics. Genome biology 20: 238.

Fagundes WC, Haueisen J, Stukenbrock EH. 2020. Dissecting the Biology of the Fungal Wheat Pathogen Zymoseptoria tritici: A Laboratory Workflow. Current protocols in microbiology 59: e128.

Faticov M, Desprez-Loustau M-L, Kiss L, Massot M, d’Arcier JF, Mutz J, Németh MZ, Roslin T, Tack AJM. 2022. Niche differentiation within a cryptic pathogen complex: climatic drivers and hyperparasitism at multiple spatial scales. Ecography 2022.

Feurtey A, Lorrain C, Croll D, Eschenbrenner C, Freitag M, Habig M, Haueisen J, Möller M, Schotanus K, Stukenbrock EH. 2020. Genome compartmentalization predates species divergence in the plant pathogen genus Zymoseptoria. BMC genomics 21: 588.

Feurtey A, Lorrain C, McDonald MC, Milgate A, Solomon PS, Warren R, Puccetti G, Scalliet G, Torriani SFF, Gout L, et al. 2023. A thousand-genome panel retraces the global spread and adaptation of a major fungal crop pathogen. Nature communications 14: 1059.

Fisher MC, Henk DA, Briggs CJ, Brownstein JS, Madoff LC, McCraw SL, Gurr SJ. 2012. Emerging fungal threats to animal, plant and ecosystem health. Nature 484: 186–194.

Flutre T, Duprat E, Feuillet C, Quesneville H. 2011. Considering Transposable Element Diversification in De Novo Annotation Approaches. PloS one 6: e16526.

Ganopoulos I, Kapazoglou A, Bosmali I, Xanthopoulou A, Nianiou-Obeidat I, Tsaftaris A, Madesis P. 2017. Application of the ITS2 region for barcoding plants of the genus Triticum L. and Aegilops L. Cereal research communications 45: 381–389.

Giraud T, Gourbière S. 2012. The tempo and modes of evolution of reproductive isolation in fungi. Heredity 109: 204–214.

Goodwin SB, M’Barek SB, Dhillon B, Wittenberg AHJ, Crane CF, Hane JK, Foster AJ, van der Lee TAJ, Grimwood J, Aerts A, et al. 2011. Finished genome of the fungal wheat pathogen Mycosphaerella graminicola reveals dispensome structure, chromosome plasticity, and stealth pathogenesis. PLoS genetics 7.

Goodwin SB, Waalwijk C, Kema GHJ, Cavaletto JR, Zhang G. 2003. Cloning and analysis of the mating-type idiomorphs from the barley pathogen Septoria passerinii. Molecular genetics and genomics: MGG 269: 1–12.

Goodwin SB, Zismann VL. 2001. Phylogenetic analyses of the ITS region of ribosomal DNA reveal that Septoria passerinii from barley is closely related to the wheat pathogen Mycosphaerella graminicola. Mycologia 93: 934–946.

Green GJ, Bendelow VM. 1961. Effect of speckled leaf blotch, Septoria passerinii Sacc., on the yield and malting quality of barley. Canadian Journal of Plant Sciences 41: 431–435.

Green GJ, Dickson JG. 1957. Pathological histology and varietal reactions in Septoria leaf blotch of barley. Phytopathology: 73–79.

Harlan JR. 1971. Agricultural origins: centers and noncenters. Science 174: 468–474.

Harlan JR, Zohary D. 1966. Distribution of wild wheats and barley. Science 153: 1074–1080.

Hoff KJ, Lomsadze A, Borodovsky M, Stanke M. 2019. Whole-Genome Annotation with BRAKER. Methods in molecular biology 1962: 65–95.

Huson DH, Bryant D. 2006. Application of phylogenetic networks in evolutionary studies. Molecular biology and evolution 23: 254–267.

Jakob SS, Blattner FR. 2006. A Chloroplast Genealogy of Hordeum (Poaceae): Long-Term Persisting Haplotypes, Incomplete Lineage Sorting, Regional Extinction, and the Consequences for Phylogenetic Inference. Molecular Biology and Evolution 23: 1602–1612.

Karisto P, Hund A, Yu K, Anderegg J, Walter A, Mascher F, McDonald BA, Mikaberidze A. 2018. Ranking Quantitative Resistance to Septoria tritici Blotch in Elite Wheat Cultivars Using Automated Image Analysis. Phytopathology 108: 568–581.

Koble AF, Peterson GA, Timian RG. 1959. A method of evaluating the reaction of barley seedlings to infection with Septoria passerinii Sacc. Plant Disease Reporter: 14–17.

Lee SH, Neate SM. 2007a. Distribution and frequency of mating types in the asexual fungus Septoria passerinii. Canadian journal of plant pathology. Revue Canadienne de phytopathologie 29: 268–275.

Lee SH, Neate SM. 2007b. Population Genetic Structure of Septoria passerinii in Northern Great Plains Barley. Phytopathology 97: 938–944.

Linde CC, Zhan J, McDonald BA. 2002. Population Structure of Mycosphaerella graminicola: From Lesions to Continents. Phytopathology 92: 946–955.

Lorrain C, Feurtey A, Möller M, Haueisen J, Stukenbrock E. 2021. Dynamics of transposable elements in recently diverged fungal pathogens: lineage-specific transposable element content and efficiency of genome defenses. G3 11.

Marks RA, Amézquita EJ, Percival S, Rougon-Cardoso A, Chibici-Revneanu C, Tebele SM, Farrant JM, Chitwood DH, VanBuren R. 2023. A critical analysis of plant science literature reveals ongoing inequities. Proceedings of the National Academy of Sciences of the United States of America 120: e2217564120.

McDonald BA, Suffert F, Bernasconi A, Mikaberidze A. 2022. How large and diverse are field populations of fungal plant pathogens? The case of Zymoseptoria tritici. Evolutionary applications 15: 1360–1373.

McNew GL. 1960. The nature, origin, and evolution of parasitism. In: Horsfall JG DAE, ed. Plant pathology: an advanced treatise. New York, NY, USA: Academic Press, 19–69.

Miller MA, Schwartz T, Pickett BE, He S, Klem EB, Scheuermann RH, Passarotti M, Kaufman S, O’Leary MA. 2015. A RESTful API for Access to Phylogenetic Tools via the CIPRES Science Gateway. Evolutionary bioinformatics online 11: 43–48.

Möller M, Habig M, Freitag M, Stukenbrock EH. 2018. Extraordinary Genome Instability and Widespread Chromosome Rearrangements During Vegetative Growth. Genetics 210: 517–529.

Monteil CL, Cai R, Liu H, Llontop MEM, Leman S, Studholme DJ, Morris CE, Vinatzer BA. 2013. Nonagricultural reservoirs contribute to emergence and evolution of Pseudomonas syringae crop pathogens. The New phytologist 199: 800–811.

Monteil CL, Yahara K, Studholme DJ, Mageiros L, Méric G, Swingle B, Morris CE, Vinatzer BA, Sheppard SK. 2016. Population-genomic insights into emergence, crop adaptation and dissemination of Pseudomonas syringae pathogens. Microbial genomics 2: e000089.

Morrell PL, Clegg MT. 2007. Genetic evidence for a second domestication of barley (Hordeum vulgare) east of the Fertile Crescent. Proceedings of the National Academy of Sciences of the United States of America 104: 3289–3294.

Nguyen L-T, Schmidt HA, von Haeseler A, Minh BQ. 2015. IQ-TREE: a fast and effective stochastic algorithm for estimating maximum-likelihood phylogenies. Molecular biology and evolution 32: 268–274.

Nishikawa T, Salomon B, Komatsuda T, von Bothmer R, Kadowaki K-I. 2002. Molecular phylogeny of the genus Hordeum using three chloroplast DNA sequences. Genome / National Research Council Canada = Genome / Conseil national de recherches Canada 45: 1157–1166.

Oggenfuss U, Badet T, Wicker T, Hartmann FE, Singh NK, Abraham L, Karisto P, Vonlanthen T, Mundt C, McDonald BA, et al. 2021. A population-level invasion by transposable elements triggers genome expansion in a fungal pathogen. eLife 10: 1–25.

Orellana-Torrejon C, Vidal T, Boixel A-L, Gélisse S, Saint-Jean S, Suffert F. 2022. Annual dynamics of Zymoseptoria tritici populations in wheat cultivar mixtures: A compromise between the efficacy and durability of a recently broken-down resistance gene? Plant pathology 71: 289–303.

Papaïx J, Burdon JJ, Zhan J, Thrall PH. 2015. Crop pathogen emergence and evolution in agro-ecological landscapes. Evolutionary applications 8: 385–402.

Penczykowski RM, Walker E, Soubeyrand S, Laine A-L. 2015. Linking winter conditions to regional disease dynamics in a wild plant-pathogen metapopulation. The New phytologist 205: 1142–1152.

Quaedvlieg W, Kema GHJ, Groenewald JZ, Verkley GJM, Seifbarghi S, Razavi M, Mirzadi Gohari A, Mehrabi R, Crous PW. 2011. Zymoseptoria gen. nov.: A new genus to accommodate Septoria-like species occurring on graminicolous hosts. Persoonia: Molecular Phylogeny and Evolution of Fungi 26: 57–69.

Quesneville H, Bergman CM, Andrieu O, Autard D, Nouaud D, Ashburner M, Anxolabehere D. 2005. Combined evidence annotation of transposable elements in genome sequences. PLoS computational biology 1: 166–175.

Quesneville H, Nouaud D, Anxolabéhère D. 2003. Detection of new transposable element families in Drosophila melanogaster and Anopheles gambiae genomes. Journal of molecular evolution 57 Suppl 1: S50–9.

Rojas-Barrera I, Fagundes C. W, Stukenbrock EH. 2023. Species of Zymoseptoria (Dothideomycetes) as a Model System to Study Plant Pathogen Genome Evolution. In: Scott B, Mesarich C, eds. The Mycota 5, Plant Relationships. Springer Nature Switzerland AG, 349–379.

Rouxel M, Mestre P, Comont G, Lehman BL, Schilder A, Delmotte F. 2013. Phylogenetic and experimental evidence for host-specialized cryptic species in a biotrophic oomycete. The New phytologist 197: 251–263.

Sang T, Crawford D, Stuessy T. 1997. Chloroplast DNA phylogeny, reticulate evolution, and biogeography of Paeonia (Paeoniaceae). American journal of botany 84: 1120.

Seifbarghi S, Razavi M, Aminian H, Zare R, Etebarian H. 2009. Studies on the host range of Septoria species on cereals and some wild grasses in Iran. Phytopathologia mediterranea 48: 422–429.

Stamatakis A. 2014. RAxML version 8: a tool for phylogenetic analysis and post-analysis of large phylogenies. Bioinformatics 30: 1312–1313.

Steenkamp ET, Wingfield BD, Desjardins AE, Marasas W. F. O. Wingfield M. J. 2002. Cryptic Speciation in Fusarium subglutinans. Mycologia 94: 1032–1043.

Stevens RB. 1960. Plant Pathology, an Advanced Treatise (J.G. Horsfall and A.E. Dimond, Ed.). Academic Press, NY.

St Pierre S, Gustus C, Steffenson B, Dill-Macky R, Smith KP. 2010. Mapping Net Form Net Blotch and Septoria Speckled Leaf Blotch Resistance Loci in Barley. Genetics and Resistance.

Stukenbrock EH. 2013. Evolution, selection and isolation: a genomic view of speciation in fungal plant pathogens. The New phytologist 199: 895–907.

Stukenbrock EH, Banke S, Javan-Nikkhah M, McDonald BA. 2007. Origin and domestication of the fungal wheat pathogen Mycosphaerella graminicola via sympatric speciation. Molecular biology and evolution 24: 398–411.

Stukenbrock EH, Bataillon T, Dutheil JY, Hansen TT, Li R, Zala M, McDonald BA, Wang J, Schierup MH. 2011. The making of a new pathogen: Insights from comparative population genomics of the domesticated wheat pathogen Mycosphaerella graminicola and its wild sister species. Genome research 21: 2157–2166.

Stukenbrock EH, Christiansen FB, Hansen TT, Dutheil JY, Schierup MH. 2012a. Fusion of two divergent fungal individuals led to the recent emergence of a unique widespread pathogen species. Proceedings of the National Academy of Sciences of the United States of America 109: 10954–10959.

Stukenbrock E, Gurr S. 2023. Address the growing urgency of fungal disease in crops. Nature 617: 31–34.

Stukenbrock EH, Quaedvlieg W, Javan-Nikhah M, Zala M, Crous PW, McDonald BA. 2012b. Zymoseptoria ardabiliae and Z. pseudotritici, two progenitor species of the septoria tritici leaf blotch fungus Z. tritici (synonym: Mycosphaerella graminicola). Mycologia 104: 1397–1407.

Tamura K, Stecher G, Kumar S. 2021. MEGA11: Molecular Evolutionary Genetics Analysis Version 11. Molecular biology and evolution 38: 3022–3027.

Thines M. 2019. An evolutionary framework for host shifts – jumping ships for survival. New Phytologist 224: 605–617.

Toubia-Rahme H, Steffenson BJ. 2004. Sources of resistance to septoria speckled leaf blotch caused by Septoria passeriniiin barley. Canadian journal of plant pathology. Revue Canadienne de phytopathologie 26: 358–364.

Treindl AD, Stapley J, Leuchtmann A. 2023. Genetic diversity and population structure of Epichloë fungal pathogens of plants in natural ecosystems. Frontiers in Ecology and Evolution 11.

Van Ginkel M, Krupinsky J, McNab A. 1999. Septoria and Stagonospora Diseases of Cereals: A Compilation of Global Research : Proceedings of the Fifth International Septoria Workshop, September 20-24, 1999, CIMMYT, Mexico. CIMMYT.

Ware SB, Verstappen CP, Breeden J, Cavaletto JR, Goodwin SB, Waalwijk C, Crous PW, Kema GHJ. 2006. Discovery of a functional Mycosphaerella teleomorph in the presumed asexual barley pathogen Septoria passerinii.

White TJ, Bruns TD, Lee SJWT, Taylor J. 1990. Amplification and direct sequencing of fungal ribosomal RNA Genes for phylogenetics. In: Innis MA, ed. PCR - Protocols and Applications - A Laboratory. 315–322.

de Wit PJGM, van der Burgt A, Ökmen B, Stergiopoulos I, Abd-Elsalam KA, Aerts AL, Bahkali AH, Beenen HG, Chettri P, Cox MP, et al. 2012. The genomes of the fungal plant pathogens Cladosporium fulvum and Dothistroma septosporum reveal adaptation to different hosts and lifestyles but also signatures of common ancestry. PLoS genetics 8: e1003088.

Yu J, Xue J-H, Zhou S-L. 2011. New universal *matK* primers for DNA barcoding angiosperms. Journal of systematics and evolution 49: 176–181.

Zhan J, Mundt CC, McDonald BA. 1998. Measuring Immigration and Sexual Reproduction in Field Populations of Mycosphaerella graminicola. Phytopathology 88: 1330–1337.

Zhan J, Pettway RE, McDonald BA. 2003. The global genetic structure of the wheat pathogen Mycosphaerella graminicola is characterized by high nuclear diversity, low mitochondrial diversity, regular recombination, and gene flow. Fungal genetics and biology: FG & B 38: 286–297.

Zheng X, Levine D, Shen J, Gogarten SM, Laurie C, Weir BS. 2012. A high-performance computing toolset for relatedness and principal component analysis of SNP data. Bioinformatics 28: 3326–3328.

