## Supplementary Information for "Emergence of sympatric host-specific lineages of the fungal plant pathogen *Zymoseptoria passerinii* in natural ecosystems"

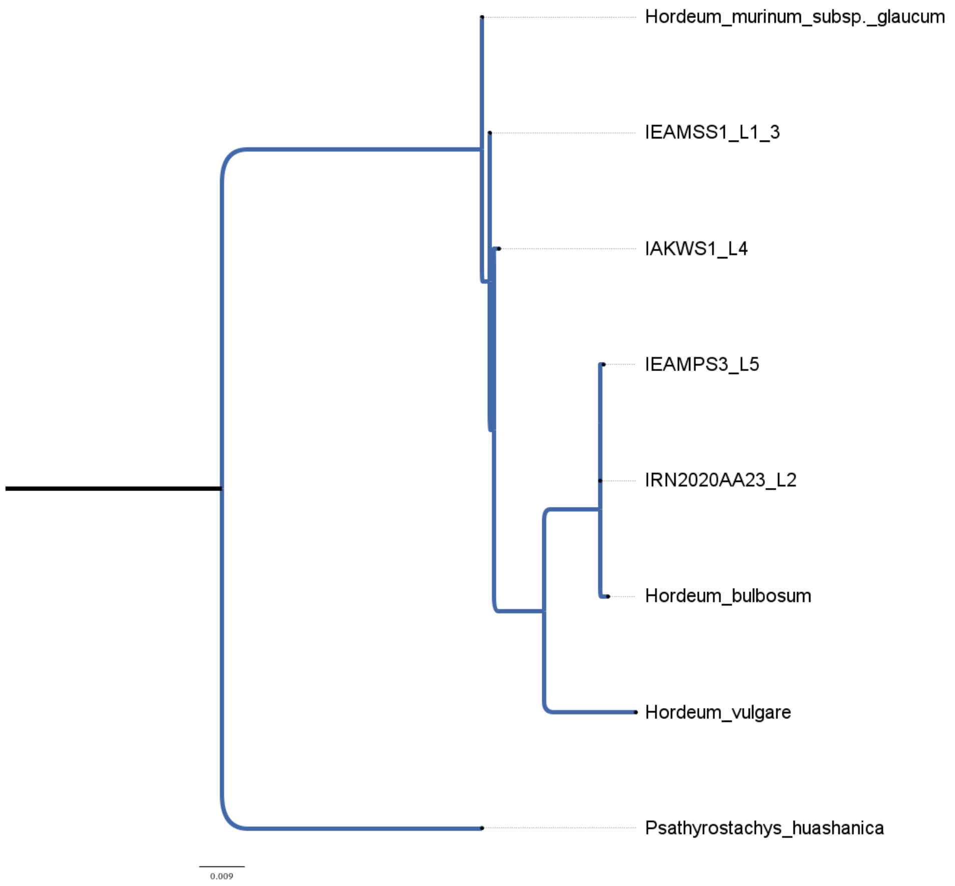


Fig S1. Maximum likelihood tree for the *Hordeum* herbarium samples that represent hosts of *Zymoseptoria passerinii*. Phylogeny with concatenated markers matK, trnH-psbA, atpB-rbcL, and trnL-trnF amplified, species identification was performed with BLAST (Table S2).

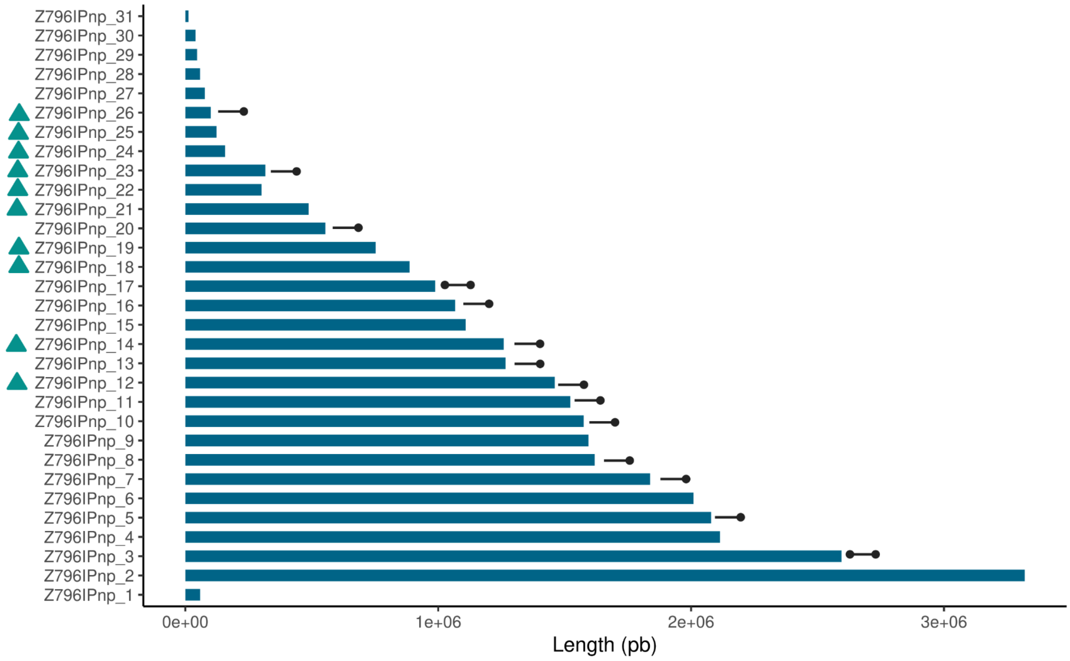


Fig. S2. Contig size for the genome assembly of the isolate Zpa796. In the accompanying bar plots, circles over the bar plots highlight contigs possessing telomeres, which may appear at one or both extremes. The ten contigs that did not align with Zpa63 genome have a triangle to the left of their ID.

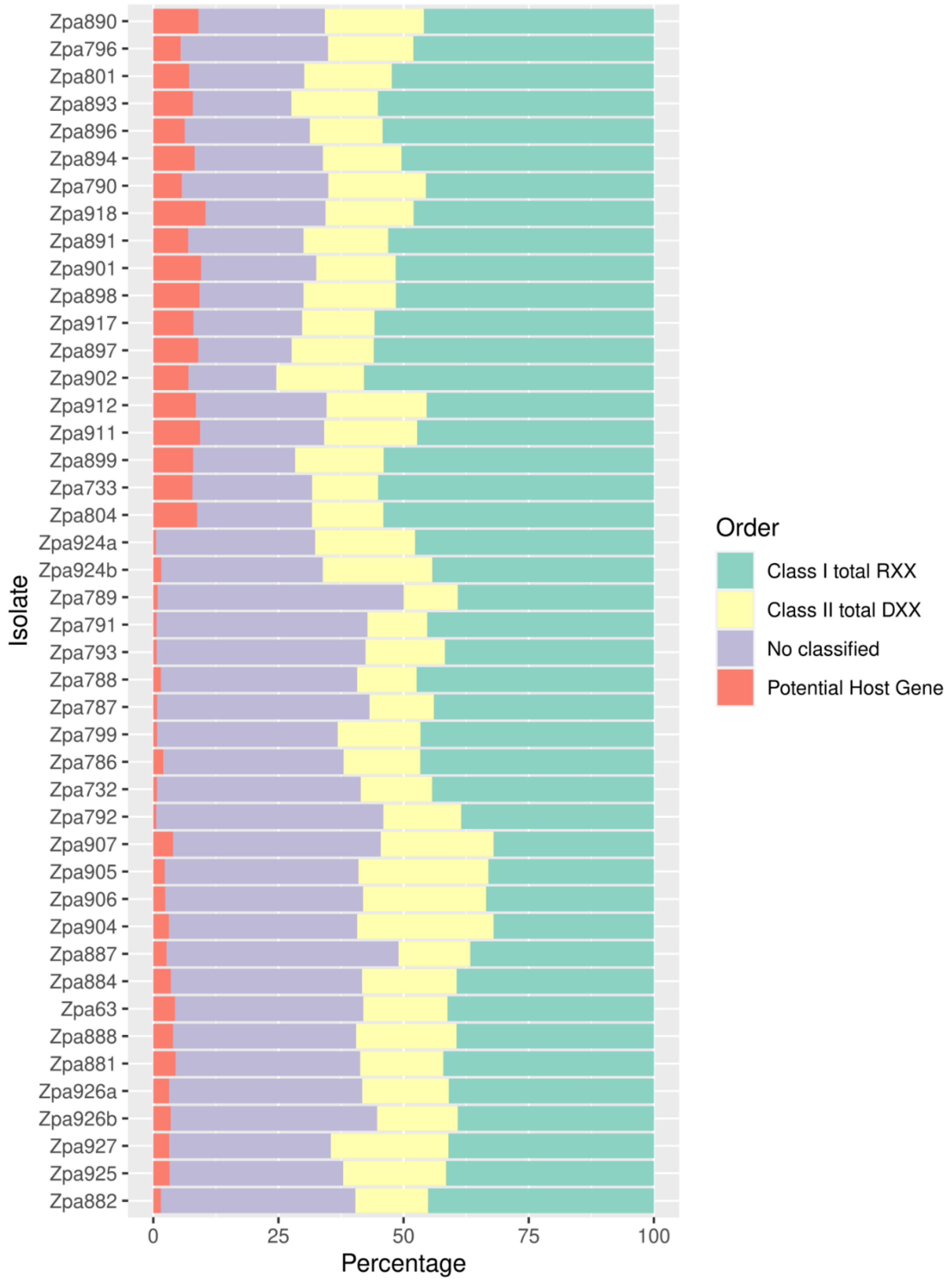


Figure S3. Proportion of repeat sequences annotated for 42 non-clonal corrected isolates of *Zymoseptoria passerinii*. Samples Zpa924a and Zpa924b; and Zpa926a and Zpa926b are technical replicates.

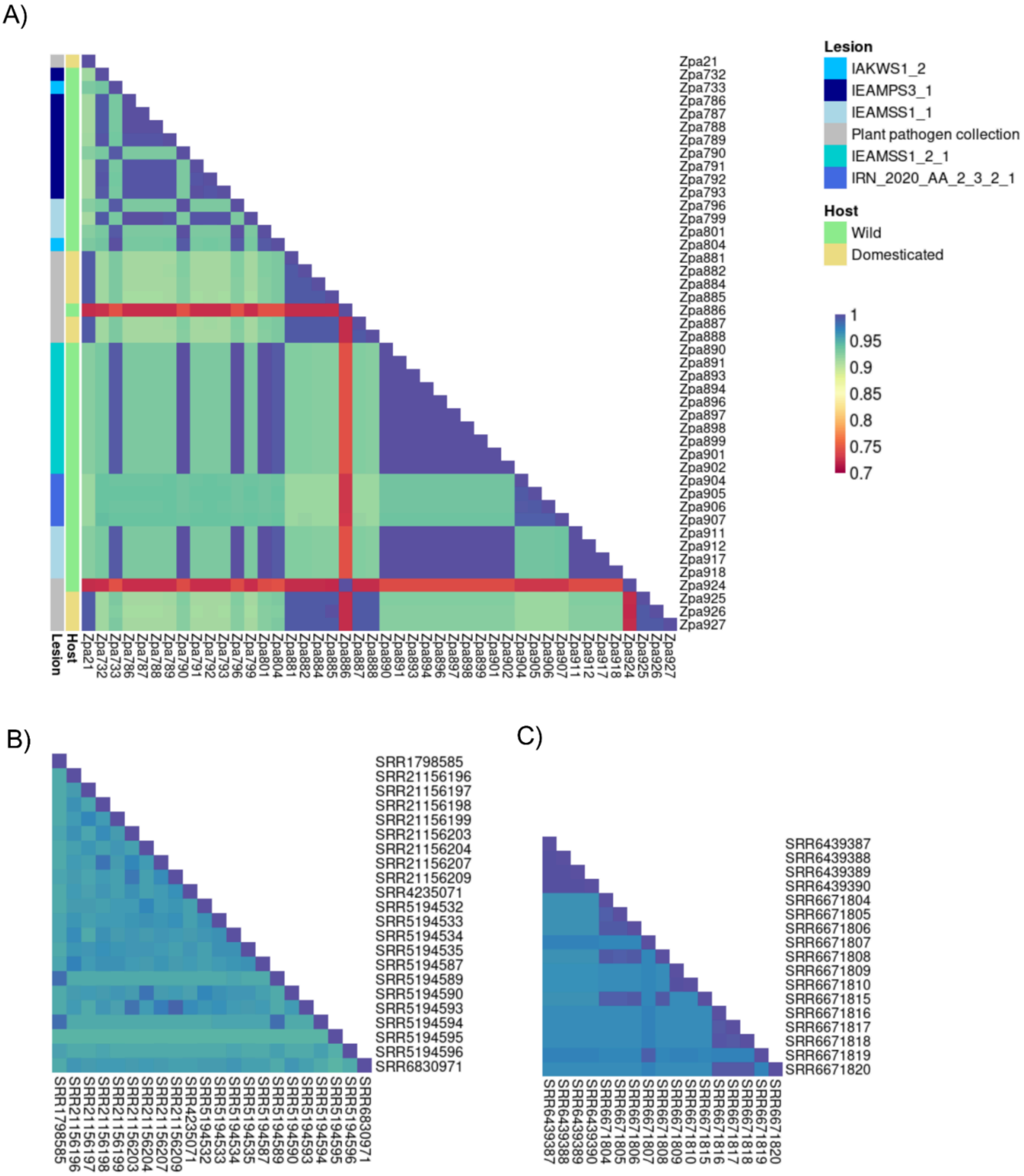


Figure S4. Identity By State (IBS) for A) *Zymoseptoria passserinii,* B) *Z. tritici, and* C) *Z. ardabiliae.* For panel A, the first vertical line shows the lesion, and the second is the type of host, domesticated (yellow) or wild grass (green). The group “Plant pathogen collection” includes all the isolates obtained from different source institutions. The color scale is valid for the three panels.

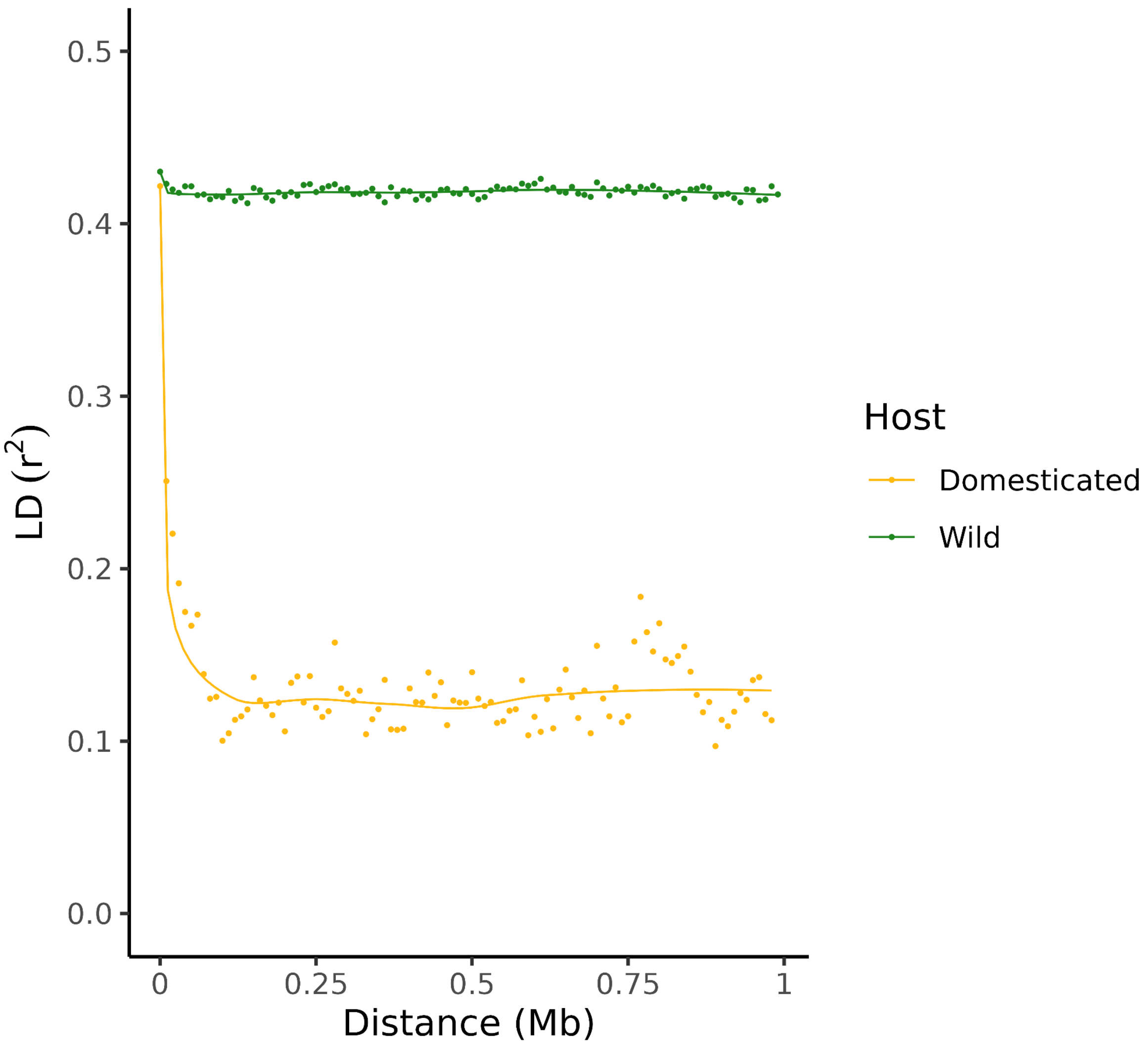


Figure S5. Linkage Disequilibrium for the contig two of *Z. passerinii*. Isolates from wild ( n = 34, green) and domesticated (n = 10, yellow) infecting hosts.

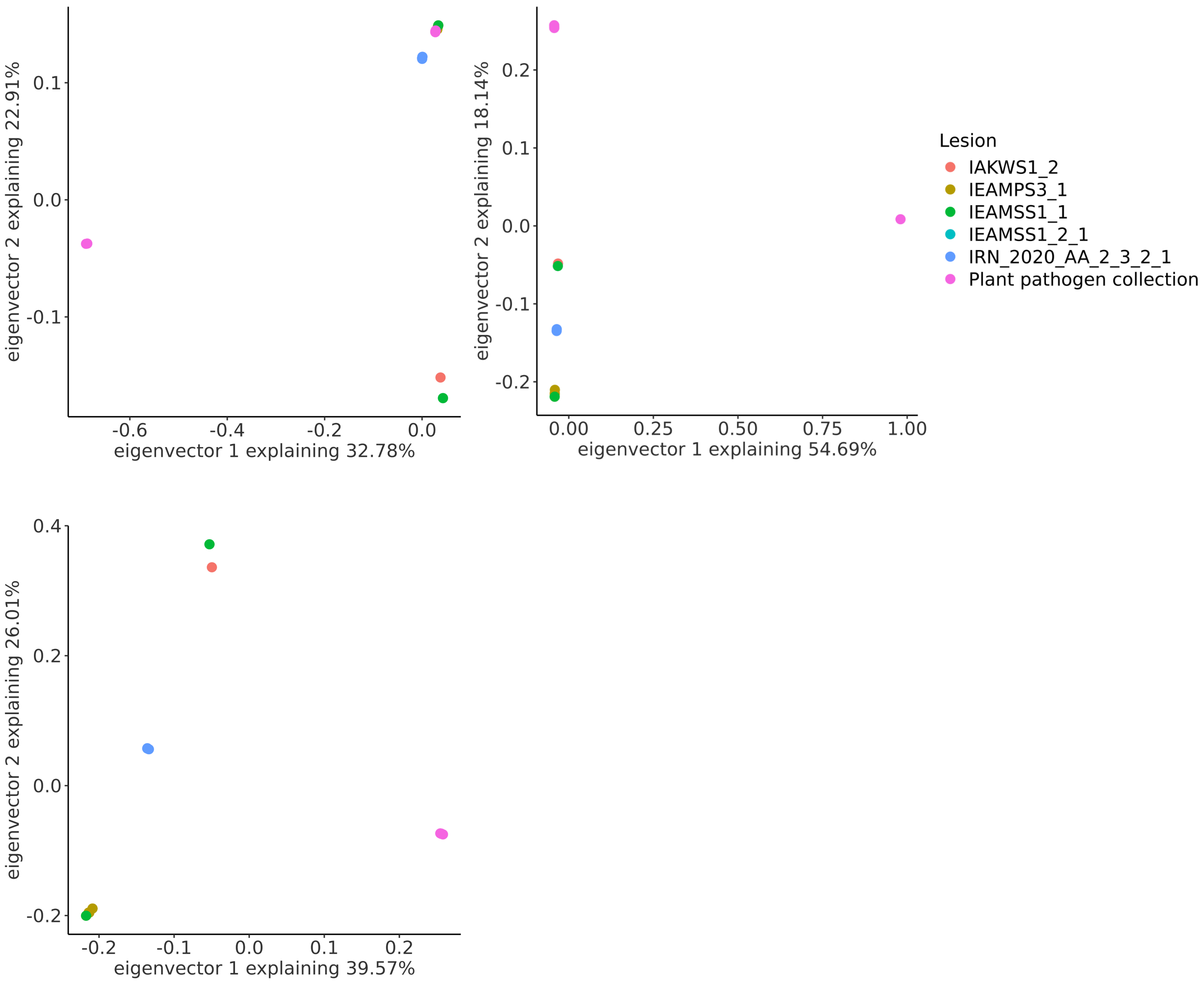


Figure S6. Principal Component Analysis plot for *Z. passerinii* a) 44 genotypes included in this study, b) non-clonal genotypes with the outlier isolate Zpa924, and c) distribution after removing Zpa924.

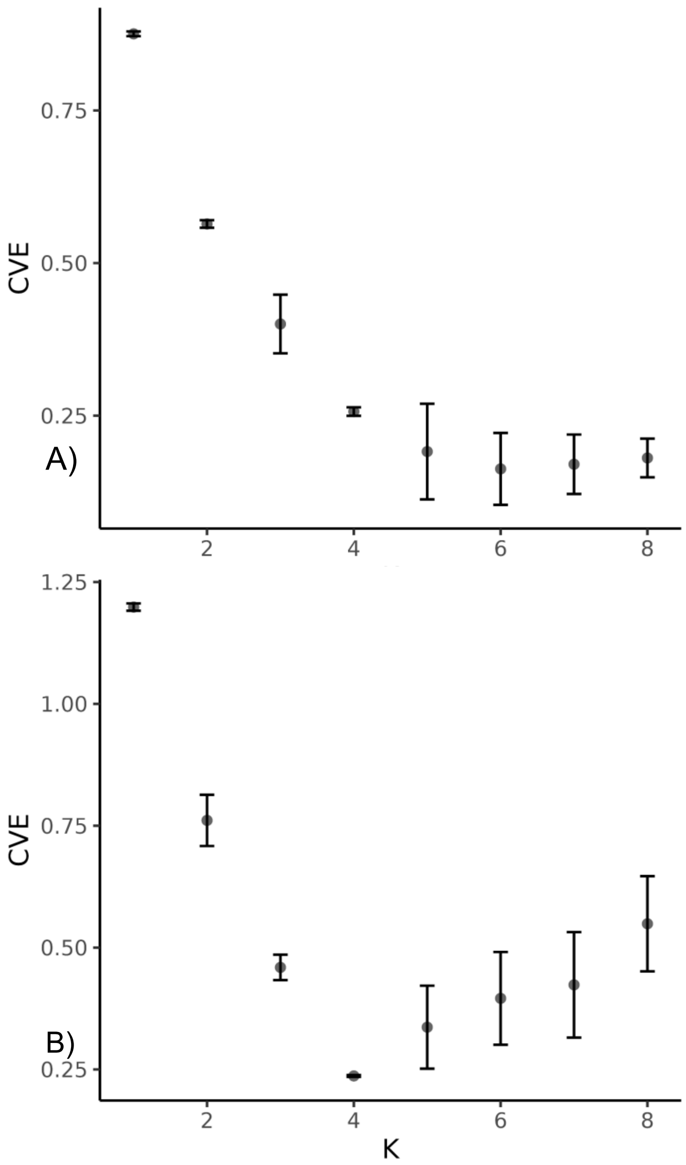


Figure S7. Mean cross-validation error (CVE) for ten admixture runs. Bars represent the standard deviation for each K-value.

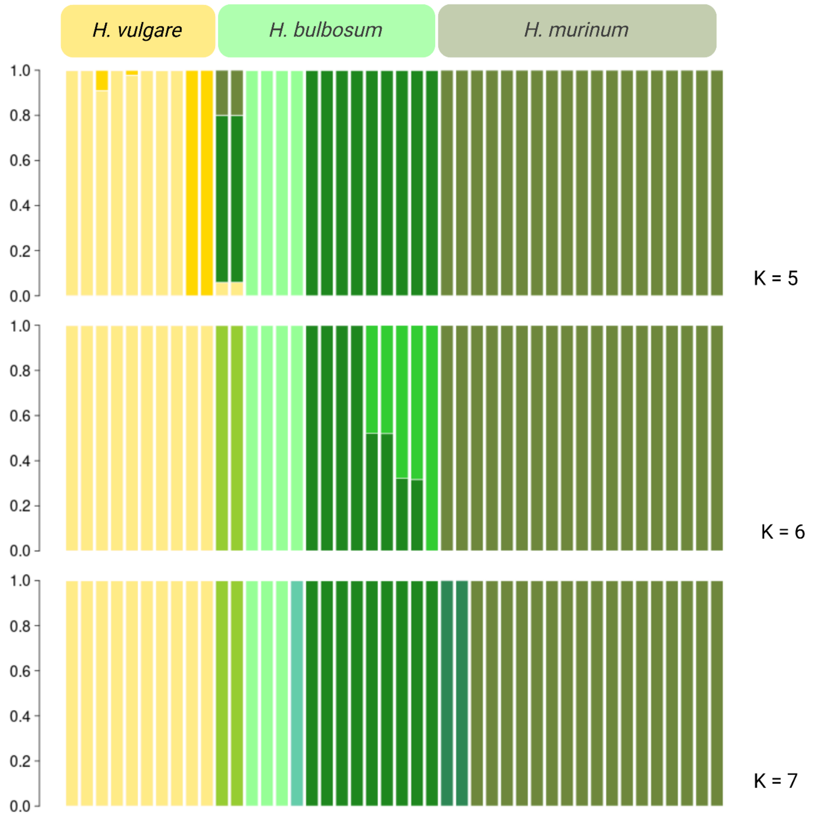


Figure S8. Ancestry analysis for the 44 genotypes (clonal and non-clonal) of *Zymoseptoria passerinii,* where the K = 6 has the lowest Cross Validation Error.

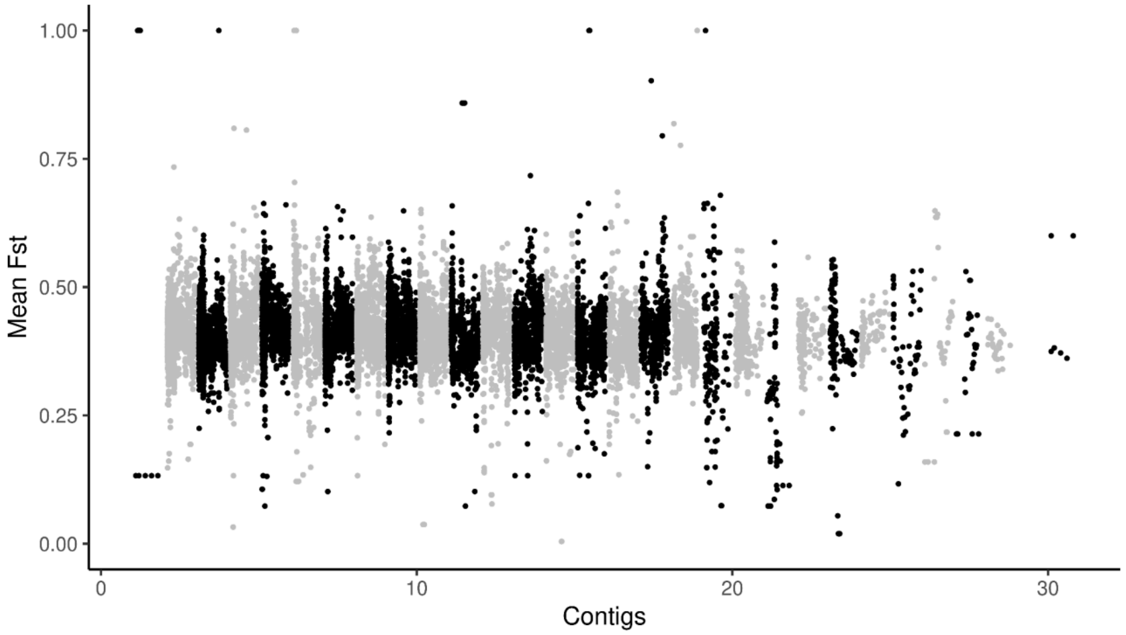


Figure S9. Whole-genome divergence ($F_{ST}$) for non-linked SNPS ($r^2 < 0.25$) between *Zymoseptoria passerinii* isolates from domesticated and wild hosts.

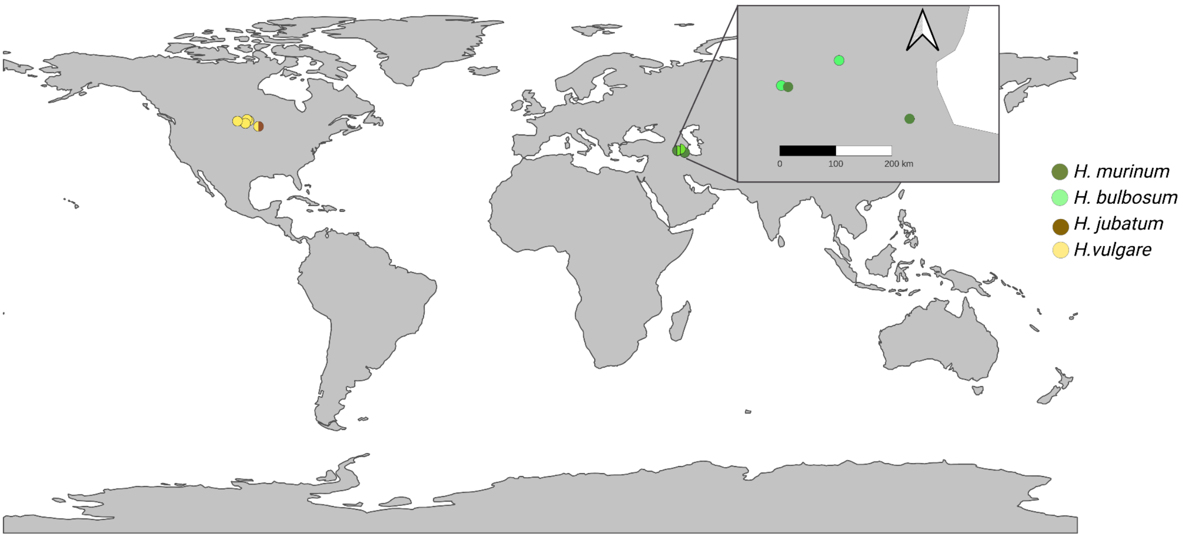


Fig. S10 Geographical location of the 42 genotypes of *Z. passerinii* included in this study.

### Material and methods

#### Genome assembly and annotation

PacBio HiFi reads from Zpa796 were assembled using the IPA HiFi Genome Assembler v1.8.0 with the --no-phase flag. Assembly metrics were obtained with Quast v5.0.2 (Gurevich *et al.*, 2013), while completeness and redundancy were evaluated with BUSCO (Manni *et al.*, 2021). The presence of fully assembled chromosomes was investigated by identifying telomeric repeats with the script FindTelomers.py (https://github.com/JanaSperschneider/FindTelomeres). Subsequently, transposable elements (TEs) were identified and annotated with the REPET pipeline v3.0 (Quesneville *et al.*, 2003, 2005; Flutre *et al.*, 2011; Ahmed *et al.*, 2011), and gene annotation was carried out with the BRAKER pipeline (Hoff *et al.*, 2019; Brůna *et al.*, 2021) using the fungal database and the transcriptomes of Zpa796 from *in vitro* cultures. Protein function was predicted with interproscan v.5.48.83 (Jones *et al.*, 2014; Blum *et al.*, 2021) with the applications SignalP_EUK-4.1 (Petersen *et al.*, 2011) and TMHMM-2.0 (Krogh *et al.*, 2001).

The Illumina reads for the 42 isolates of *Z. passerinii* and the two technical replicates were assembled with SPAdes v3.11.1 (Bankevich *et al.*, 2012), and assembly statistics were computed with Quast v5.0.2 (Gurevich *et al.*, 2013). Contigs smaller than 1 Kb were filtered out, and gene annotation was performed with BRAKER (Hoff *et al.*, 2019; Brůna *et al.*, 2021) using the gff obtained from the Zpa796 PacBio assembly. TEs were identified with the REPET pipeline v2.5 (Quesneville *et al.*, 2003, 2005; Flutre *et al.*, 2011; Ahmed *et al.*, 2011).

#### Single Nucleotide Variant pipeline for Zymoseptoria species

The Zpa796 PacBio assembly was used to perform the Single Nucleotide Variants (SNVs) calling for a total of 93 Illumina genomes: 44 of *Z. passerinii,* 21 of *Z. ardabiliae*, 22 of *Z. tritici*, 4 for *Z. pseudotritici*, 2 for *Z. brevis*. The SRA and Bioproject numbers of genome downloaded from the NCBI are listed in Table S4.

The paired fastq files were trimmed with bbduk (Bushnell, 2016) with the parameters: trimpolya=10 ktrim=r k=23 mink=11 hdist=1 qtrim=rl trimq=20 maq=20 tpe tbo; the quality of the sequences was assessed with FastQC (Andrews, 2010) and a summary report was built multiqc (Ewels *et al.*, 2016). The trimmed sequences were aligned against the genome assembly of Zpa796 with NextGenMap 0.5.5 (Sedlazeck *et al.*, 2013). The SAM files were sorted with samtools 1.9 (Danecek *et al.*, 2021), and the picard options AddOrReplaceReadGroups and MarkDuplicates, were used to edit the read groups information and mark duplicates, respectively. The SNVs calling was performed following the pipeline of Genome Analysis Toolkit (GATK), first gvcf files were generated with HaplotypeCaller -ploidy 1 -ERC GVCF, and these were combined with CombineGVCFs using the options --read-filter MappingQualityReadFilter --read-filter OverclippedReadFilter. The genotyping was performed with GenotypeGVCFs option, a total of 6,549,866 non-filter SNVs were called. Filtering was performed in multiple steps and per species. The Variant Call Format (VCF) file obtained with GATK was first filtered with vcftools v 0.1.14 with the parameters --max-missing 0.5 --mac 3 --minQ 30. Subsequently, missingness per individual was computed with --missing-indv flag, and those genotypes with missing data higher than 0.1 were discarded. Missingness per site was calculated per species with the flag --missing-site; those sites with more than 0.1 missing data were removed. As describe in the main text, efter filtering the VCF file, we kept 65 genotypes and 1,167,977 SNVs for the five species of *Zymoseptoria*.

#### Clone correction and exploratory analysis

We sequenced two technical replicates for two different genotypes (Zpa924a-b and Zpa926a-b) (Table S3). IBS values were computed for the five species using PLINK v1.90b6.26 with the flag --distance 'ibs' (Chang *et al.*, 2015). The technical replicates had an IBS equal to 0.999; therefore, if two or more isolates from the same leaf had an IBS equal to or higher ≥ 0.999, these were considered clonal genotypes, an IBS value below this threshold was considered as biological variation (Figure S4). After clone correction, we kept 65 genotypes out of 93: 26 out of 42 isolates of *Z. passerinni*, 14 out of 21 for *Z. aradabiliae*, one out of two for *Z. brevis*, two out of four for *Z. pseudotritici*, and 22 out of 22 for *Z. tritici*. Linkage disequilibrium was calculated with PLINK v1.90b6.2 with the flags --r2 --ld-window-bp 50000; genotypes were grouped per type of host: domesticated and wild *Hordeum* (Figure S5).

#### Inoculation of *Z. passerinii* in *Hordeum*

For the inoculation, cells were harvested, and the fungal inoculum was adjusted to 1x106 cells/mL for Zpa63; since Zpa796 predominantly has a hyphal growth in liquid culture and tends to melanize at high cell density, the inoculum was adjusted to 2.5 x105 hyphye/mL; in both cases, cells were inoculated in a solution of Tween 20 (Roth, Karlsruhe, Germany) 0.1% [v/v], the mock treatment consisted of the Tween solution without fungal cells. We inoculated the second leaf of *Hordeum* seedlings, which emerged on day 14 after germination, with *H. vulgare* ssp. *vulgare*, *H. jubatum*, and *H. vulgare* ssp. *spontaneum* (HOR2680), and on days 14-16 for *H. murinum* ssp. *glaucum* (GRA3223). The leaves were inoculated using a spray gun and then enclosed in plastic bags for 48 hours to create an environment with maximum relative humidity.

#### References


**Ahmed I, Sarazin A, Bowler C, Colot V, Quesneville H**. **2011**. Genome-wide evidence for local DNA methylation spreading from small RNA-targeted sequences in Arabidopsis. *Nucleic acids research* **39**: 6919–6931.

**Andrews S**. **2010**. FastQC A Quality Control tool for High Throughput Sequence Data.

**Bankevich A, Nurk S, Antipov D, Gurevich AA, Dvorkin M, Kulikov AS, Lesin VM, Nikolenko SI, Pham S, Prjibelski AD, *et al.*** **2012**. SPAdes: a new genome assembly algorithm and its applications to single-cell sequencing. *Journal of computational biology: a journal of computational molecular cell biology* **19**: 455–477.

**Blum M, Chang H-Y, Chuguransky S, Grego T, Kandasaamy S, Mitchell A, Nuka G, Paysan-Lafosse T, Qureshi M, Raj S, *et al.*** **2021**. The InterPro protein families and domains database: 20 years on. *Nucleic acids research* **49**: D344–D354.

**Brůna T, Hoff KJ, Lomsadze A, Stanke M, Borodovsky M**. **2021**. BRAKER2: automatic eukaryotic genome annotation with GeneMark-EP+ and AUGUSTUS supported by a protein database. *NAR genomics and bioinformatics* **3**: lqaa108.

**Bushnell B**. **2016**. BBTools. *DOE Joint Genome Institute*.

**Chang CC, Chow CC, Tellier LC, Vattikuti S, Purcell SM, Lee JJ**. **2015**. Second-generation PLINK: rising to the challenge of larger and richer datasets. *GigaScience* **4**: 7.

**Danecek P, Bonfield JK, Liddle J, Marshall J, Ohan V, Pollard MO, Whitwham A, Keane T, McCarthy SA, Davies RM, *et al.*** **2021**. Twelve years of SAMtools and BCFtools. *GigaScience* **10**.

**Ewels P, Magnusson M, Lundin S, Käller M**. **2016**. MultiQC: summarize analysis results for multiple tools and samples in a single report. *Bioinformatics* **32**: 3047–3048.

**Flutre T, Duprat E, Feuillet C, Quesneville H**. **2011**. Considering Transposable Element Diversification in De Novo Annotation Approaches. *PloS one* **6**: e16526.

**Gurevich A, Saveliev V, Vyahhi N, Tesler G**. **2013**. QUAST: quality assessment tool for genome assemblies. *Bioinformatics* **29**: 1072–1075.

**Hoff KJ, Lomsadze A, Borodovsky M, Stanke M**. **2019**. Whole-Genome Annotation with BRAKER. *Methods in molecular biology* **1962**: 65–95.

**Jones P, Binns D, Chang H-Y, Fraser M, Li W, McAnulla C, McWilliam H, Maslen J, Mitchell A, Nuka G, *et al.*** **2014**. InterProScan 5: genome-scale protein function classification. *Bioinformatics* **30**: 1236–1240.

**Krogh A, Larsson B, von Heijne G, Sonnhammer EL**. **2001**. Predicting transmembrane protein topology with a hidden Markov model: application to complete genomes. *Journal of molecular biology* **305**: 567–580.

**Manni M, Berkeley MR, Seppey M, Simão FA, Zdobnov EM**. **2021**. BUSCO Update: Novel and Streamlined Workflows along with Broader and Deeper Phylogenetic Coverage for Scoring of Eukaryotic, Prokaryotic, and Viral Genomes. *Molecular biology and evolution* **38**: 4647–4654.

**Petersen TN, Brunak S, von Heijne G, Nielsen H**. **2011**. SignalP 4.0: discriminating signal peptides from transmembrane regions. *Nature methods* **8**: 785–786.

**Quesneville H, Bergman CM, Andrieu O, Autard D, Nouaud D, Ashburner M, Anxolabehere D**. **2005**. Combined evidence annotation of transposable elements in genome sequences. *PLoS computational biology* **1**: 166–175.

**Quesneville H, Nouaud D, Anxolabéhère D**. **2003**. Detection of new transposable element families in Drosophila melanogaster and Anopheles gambiae genomes. *Journal of molecular evolution* **57 Suppl 1**: S50–9.

**Sedlazeck FJ, Rescheneder P, von Haeseler A**. **2013**. NextGenMap: fast and accurate read mapping in highly polymorphic genomes. *Bioinformatics* **29**: 2790–2791.
